# *FvTFL1* reverses the function of *FvGI-FvCO-FvFT1* pathway in the photoperiodic flowering of woodland strawberry

**DOI:** 10.64898/2026.04.30.721829

**Authors:** Quan Zhou, Sergei Lembinen, Tuomas Toivainen, Takeshi Kurokura, Guangxun Fan, Paula Elomaa, Elli Koskela, Timo Hytönen

## Abstract

- Photoperiod is a stable seasonal signal. Although the photoperiodic flowering is well understood in short-day (SD) and long-day (LD) annual plants, regulatory mechanisms in perennials remain elusive. In a perennial woodland strawberry (*Fragaria vesca* L.), flowering is induced in SDs in autumn and plants flower following spring, while in plants with mutated *FvTERMINAL FLOWER1* (*FvTFL1*), LDs induce flowering.
- We investigated photoperiodic flowering of *F. vesca* through phenotypic and molecular characterization of transgenic lines and their crosses. We studied natural variation in flowering time and gene expression in European accessions, and explored their correlations with climatic, geographical and genetic origins.
- We showed that *FvGIGANTEA* (*FvGI*) and *FvCONSTANS* (*FvCO*) activate *FvFLOWERING LOCUS T1* (*FvFT1*) in LDs resulting in early flowering in *fvtfl1* mutant, while in SD *F. vesca*, activation of *FvTFL1* by FvFT1 reverses the photoperiodic requirement of flowering. In natural accessions, decreasing expression of *FvFT1* and *FvTFL1* towards colder climates in the east and north correlated with earlier flowering.
- We define a photoperiodic flowering mechanism controlling floral transition of perennial *F. vesca* in autumn that differs from known mechanisms in annual and perennial plants. Our findings open new avenues to understand how perennial plants cope with changing seasons across climatic and geographical ranges.

## INTRODUCTION

Perennials comprise the largest fraction of plant species on Earth with an increasing proportion towards higher latitudes (Poppenwimer *et al*., 2023). They are mostly polycarpic plants that repeat yearly cycles of vegetative and generative growth to ensure successful reproduction during their lifespan, in contrast to monocarpic annual and biennial species that rely on a single flowering event (Albani & Coupland, 2010). A common perennial life cycle comprises of formation of overwintering floral buds in autumn as a response to short days (SD) and/or decreasing temperatures and flowering in the following spring (Schnablová *et al*., 2021). Such life cycle is regulated by strong floral repressors that restrict floral transition to certain meristems and time window, conferring seasonal flowering and the maintenance of vegetative meristems for the next yearly cycle (Wang *et al*., 2009; Albani & Coupland, 2010; Wang *et al*., 2011; Koskela *et al*., 2012; Lazaro *et al*., 2018). Although photoperiod is a major signal controlling seasonal flowering, underlying mechanisms are poorly understood in perennials (Albani & Coupland, 2010; Andrés & Coupland, 2012).

In perennial woodland strawberry (*Fragaria vesca* L.), *TERMINAL FLOWER1* (*FvTFL1*) encodes a floral repressor that confers seasonal flowering habit (Koskela *et al*., 2012). The peculiarity of *F. vesca* is that the presence of functional *FvTFL1* reverses the photoperiodic requirement for flower induction. Wild type plants require SDs in autumn for flower induction in the temperature range of 13-20°C and flower seasonally, whereas *fvtfl1* mutants are quantitative long day (LD) plants that flower perpetually (Heide & Sønsteby, 2007; Koskela *et al*., 2012; Rantanen *et al*., 2015). TFL1 belongs to the family of phosphatidylethanolamine binding proteins (PEBP), that delay floral transition, promote the indeterminacy of inflorescence meristem and contribute to perennial life cycle by blocking flowering of young axillary shoots (Bradley *et al*., 1996; Bradley *et al*., 1997; Wang *et al*., 2011; Lembinen *et al*., 2023). A gene encoding another PEBP, *FLOWERING LOCUS T* (*FT*) is expressed under flower-inductive conditions in both LD and SD plants, and FT protein functions as a mobile flower-inducing signal, the florigen (Kobayashi *et al*., 1999; Faure *et al*., 2007; Tamaki e*t al.*, 2007; Komiya *et al*., 2009; Kong *et al*., 2010). FT is synthesized in leaves and transported to the shoot apical meristem (SAM), where it interacts with the bZIP transcription factor FD to induce flowering (Abe *et al*., 2005). In the SAM, FT has been shown to compete with TFL1 for binding with FD (Zhu *et al*., 2020), providing a possible mechanism for their antagonistic functions. However, in *F. vesca*, expression patterns of *FT*-homologs suggest different mechanism in the regulation of photoperiodic flower induction. *FvFT1* is expressed in the leaves specifically in LDs in both wild type and *fvtfl1* mutant plants, correlating positively with flowering only in the mutant (Koskela *et al*., 2012; Rantanen *et al*., 2014; Kurokura *et al*., 2017), while *FvFT2* and *FvFT3* are highly activated in the late inflorescence and flower meristems (Koskela *et al*., 2017; Lembinen *et al*., 2023).

Homologs of GIGANTEA (GI) and CONSTANS (CO) are major activators of florigens in both SD and LD annual plants (Song *et al*., 2015; Vicentini *et al*., 2023). In *Arabidopsis*, GI interacts with the blue-light receptor FLAVIN-BINDING, KELCH REPEAT, F-BOX 1 (FKF1) to promote *CO* transcription in LDs (Imaizumi *et al*., 2003; Sawa *et al*., 2007). GI and FKF1 also contribute to the stabilization of CO protein that accumulates when *CO* mRNA expression coincides with light in the afternoon, enabling CO to activate *FT* in leaves (Valverde *et al*., 2004; Hwang *et al*., 2019). A recent work showed that *CO-FT/TWIN SISTER OF FT-LIKE (TSFL)* module promotes flowering in LDs also in the *perpetual flowering1* (*pep1*) mutant of alpine rockcress (*Arabis alpina* L.), a perennial relative of Arabidopsis. Both *AaCO* and *AaFT/TSFL* genes are involved in the regulation of flowering in the primary shoot, while *AaCO* is not needed for flowering in axillary shoots, indicating that the photoperiodic pathway contributes also to perennial shoot architecture (Sashidhar & Coupland, 2026). Similarly, *FvCO* and *FvFT1* were shown to promote flowering in LDs in *F. vesca fvtfl1* mutant, and *FvCO* was found to be essential for the activation of *FvFT1* (Koskela *et al*., 2012; Rantanen *et al*., 2014; Kurokura *et al*., 2017). Because these studies in *A. alpina* and *F. vesca* were carried out in perpetual flowering mutants in which LDs promote flowering, they do not provide answers to a major open question: How does photoperiod regulate floral transition in perennials in autumn?

Using transgenic lines and their crosses, we show that LDs activate *FvFT1* through conserved *FvGI-FvCO-FvFT1* photoperiodic flowering pathway in both seasonal flowering SD genotypes and perpetual flowering LD genotypes of *F. vesca*. We provide direct evidence that the opposite photoperiodic responses of these genotypes arise from epistatic interactions between *FvFT1* and *FvTFL1*; FvFT1 upregulates *FvTFL1* in LDs to repress flowering in seasonal flowering accessions, while in the absence of *FvTFL1*, FvFT1 activates *FvAP1* and flowering. Further analysis on European *F. vesca* accessions, that are divided into eastern and western genetic clusters (Toivainen *et al*., 2026), revealed lower average *FvFT1* expression level and earlier flowering in eastern *F. vesca* populations compared with western accessions. Also, a strong positive correlation between *FvFT1* and *FvTFL1* mRNA levels was found, indicating that *FvFT1* is a major driver of geographical differences in flowering time. Overall, our findings define a photoperiodic flowering mechanism in strawberries that differs from those described in annual and perennial plants and offers potential strategies to optimize flowering and yield under changing climate in strawberries and related fruit crops of the Rosaceae family.

## MATERIALS AND METHODS

### Plant material

*Fragaria vesca* accessions Hawaii-4 (H4, *fvtfl1* mutant), FIN56 (PI551792, originally obtained from the National Clonal Germplasm Repository, Corvallis) and NOR8 from Alta, Norway (70,03244°N, 23,40124°E) were maintained as seeds through self-pollinations and used for gene functional analyses. FIN56 and NOR8 were previously fully sequenced and showed low genome-wide heterozygosity levels (Toivainen *et al*., 2026). Newly generated H4-*FvGI*-RNAi and H4-*FvGI* overexpression (OE) lines, and previously reported H4-*FvFT1*-RNAi and H4-*FvCO*-RNAi lines were germinated from seeds and used for gene functional studies (Koskela *et al*., 2012; Kurokura *et al*., 2017). *FvGI*-OE lines were generated also in seasonal flowering FIN56 and NOR8 backgrounds. These materials were vegetatively propagated for experiments through runners. To investigate genetic interactions among *FvGI, FvCO, FvFT1* and *FvTFL1*, several crosses between the transgenic lines and selected accessions were carried out and resulting F1 and F2 hybrids were used for the experiments (Table **S1)**. Finally, a collection of 181 European *F. vesca* accessions, maintained as clones, was used for studying natural variation in flowering time (Table **S2**).

### Plasmid constructs and transformation

Plasmid constructs for overexpression lines were created according to Gateway technology with Clonase II (Invitrogen). For *FvGI-OE* and *FvGI*-RNAi lines, cDNA sequences from *F. vesca* accession H4 were amplified using primers described in Table **S3** and introduced into the pH7WG2D-1 (overexpression) and pK7GWIWG2-7F2 (RNAi-silencing) destination vectors containing GFP as a positive selection marker (Karimi *et al*., 2002). Strawberry transformation was carried out as described previously (Oosumi *et al*., 2006). 13-20 transgenic lines were generated and screened for flowering time phenotypes and changes in gene expression for each construct. 3-4 lines per construct were selected for further experiments.

### Growth conditions and phenotyping

H4 and its transgenic lines were germinated from seeds on wet filter paper in 18-h LDs (LD^18h^) at 22 °C until cotyledons were visible. At the two-cotyledon stage, seedlings were rooted to Jiffy Peat Disks (Jiffy Products International BV) in a greenhouse in LD^18h^ at 20/17 °C (day/night). After 2 weeks, plants with 2-3 leaves were transplanted into 8 × 8 cm pots for photoperiodic treatments. Photoperiodic treatments were conducted in a greenhouse under LD^18h^ (18h light/6h dark) or SD^12h^ (12h light/12h dark) conditions. In the greenhouse, natural lighting was complemented with 150 µmol m–2 s–1 PPFD from high-pressure sodium lamps (Airam 400W, Kerava, Finland).

Runner plants of NOR8, FIN56 and their transgenic lines were rooted on Jiffy peat disks (Jiffy Products International BV) and then transferred into 8cm x 8cm pots filled with peat (Kekkilä, Finland). The plants with at least 2–3 leaves were subjected to photoperiodic treatments indicated in the figure legends in either greenhouse (FIN56) or growth rooms (NOR8) at 17°C. Growth rooms were equipped with LED lamps (AP67, Valoya, Finland), providing a photosynthetic photon flux density (PPFD) of ∼150 µmol m⁻² s⁻¹. After the treatments, plants were transferred to the greenhouse for flowering time observations using the conditions described above.

All hybrids were propagated in the greenhouse (LD^18h^ at 20/17 °C), and plants were subjected to photoperiodic treatments in either growth rooms described above or in Arabidopsis Chamber AR-36L2 (Percival, Germany), followed by flowering observations in the greenhouse (LD^18h^ at 20/17 °C). Conditions during the photoperiodic treatments are indicated in figure legends.

To explore flowering time of natural accessions, the plants were subjected to SD^12h^ at 16°C for six weeks in the greenhouse. In another experiment, the accessions were exposed to ambient photoperiod and temperature conditions in Helsinki, Finland, from mid-August until October 12, 2017 (Field Aug-Oct). Both groups were transferred to greenhouse (LD^18h^, 20/17°C) after treatments for flowering observations. All flowering-time phenotypes were determined as the number of leaves initiated before the first inflorescence and the number of days from the beginning/end of the photoperiodic treatments (indicated in figure legends) to the first flower open.

### RNA extraction and gene expression analyses

Leaf and shoot apex samples were collected under photoperiodic treatments in time points indicated in the figures and stored at −80°C before carrying out RNA extraction. Each biological replicate consisted of a single leaflet or one or three shoot apices. Total RNA was extracted from leaf and apex samples as in Mouhu *et al*. (2009). Protoscript II reverse transcriptase kit (NEB, US) was used for cDNA synthesis. qRT-PCR was performed using a LightCycler 480 SYBR Green I Master kit (Roche Diagnostics, 555 Indianapolis, US) in a Roche LightCycler 480 (Roche Diagnostics, Indianapolis US), as described in Mouhu *et al*., (2009). Primers used in this study are shown in Table **S3**. Each gene was tested by three technical replicates. *FvMSI1 (FvH4_7g08380*) was used as a stable reference gene (Mouhu *et al*., 2009). The number of biological replicates was described in the figure legends.

### RNA sequencing

For bulk RNA-seq, total RNA was extracted from leaves collected at ZT4 from three biological replicates of 12 accessions (six western and six eastern; Dataset **S1**) reported in Toivainen *et al*., (2026), grown in a greenhouse under LD^18h^ at 18 °C. Libraries were prepared using the TruSeq Total RNA kit (Illumina) following rRNA depletion with Pan-Plant riboPOOL probes (siTOOLs Biotech) and MagSi-STA 1.0 beads (Magtivio). Sequencing (100 bp, single-end) was performed on an Illumina NextSeq500. Raw FASTQ reads were quality-trimmed with Trimmomatic v0.39 in single-end mode (SLIDINGWINDOW:5:20, LEADING:5, TRAILING:5, MINLEN:50). Trimmed reads were aligned to the *Fragaria vesca* reference genome v.4.0 using STAR v2.7.10a with the genome indexed using the v4.0.a2 annotation from the Genome Database for Rosaceae. Only uniquely mapped reads were retained. Gene-level counts were generated with HTSeq-count v0.13.5 in unstranded mode.

### Scanning electron microscopy

Dissected meristem samples were fixed overnight in FAA buffer (3.7% formaldehyde, 5% acetic acid, and 50% ethanol) and then passed through an ethanol series (50%, 60%, 70%, 80%, 90%, and 100%) under mild vacuum (∼0.6 atm). Critical point drying was performed using a Leica EM CPD300 (Leica Mikrosystems GmbH, Vienna, Austria). The dried samples were coated with platinum using a Quorum Q150TS coater (Quorum Technologies, UK) and examined using a Quanta 250 FEG scanning electron microscope (FEI, Hillsboro, OR, USA) at the Electron Microscopy Unit, Institute of Biotechnology, University of Helsinki.

### Statistical analyses

For flowering time data from H4, NOR8, their transgenic lines, and F1 hybrids described in Fig. **3a**, mean values were analyzed by one-way ANOVA followed by Tukey’s HSD multiple comparisons. Flowering time data from *F. vesca* European accessions, other F1 hybrids (Fig. **4a**), and F2 populations (Fig. **4e, f**) were analyzed by ANOVA followed by pairwise t-tests with Tukey’s HSD. Pearson correlation coefficients were calculated between flowering time and relative gene expression, and among the expression levels of flowering-time genes. P-values were estimated with Bonferroni correction. All analyses were performed in R v4.3.0 (R Core Team 2023).

### Sequence alignment and phylogenetic analysis

Protein sequences were retrieved from the NCBI database (Table **S4**). Multiple sequence alignment was performed in MEGA11 using the ClustalW algorithm with default gap opening and extension penalties. The alignment was inspected manually to confirm correct positional homology and adjusted where necessary. Phylogenetic analysis was conducted in MEGA11 using the Neighbor-Joining method. Branch support was assessed with 1,000 bootstrap replicates, and gaps/missing data were treated using the pairwise-deletion option. Sequence conservation among the six species, which GI homolog function has been reported, was evaluated using DNAman v8.0. Conserved regions were identified using the DNAman conservation analysis module, and percent identity was calculated across the full-length alignment as well as within key conserved blocks which were shown in different colors.

## RESULTS

### *FvGI*, *FvCO* and *FvFT1* promote flowering in long days in perpetual flowering *F. vesca tfl1* mutant

We have previously identified key regulators of photoperiodic pathway in *F. vesca* including *FvCO* (FvH4_6g45860; Kurokura *et al*., 2017) and *FvFT1* (FvH4_6g00090; Koskela *et al*., 2012). Here, we analysed a single gene encoding a GI-like protein from *Fragaria vesca* genome v4.0 (FvH4_2g13030; Li *et al*., 2019). FvGI is highly conserved with GI homologues from other species and is positioned among the GI homologs from other Rosaceous species in the phylogenetic tree (Fig. **S1**). To explore the role of *FvGI* in the photoperiodic flowering pathway of *F. vesca,* we generated transgenic *FvGI* overexpression (*FvGI*-OE) and RNAi silencing lines in the perpetual flowering *fvtfl1* mutant accession H4 and examined their flowering phenotypes under LD^18h^ and SD^12h^ conditions together with previously developed *FvCO*-RNAi and *FvFT1*-RNAi silencing lines (Koskela *et al*., 2012, Kurokura *et al*., 2017, Fig. **S2A-D**). Similarly to *FvCO*-RNAi and *FvFT1*-RNAi plants, *FvGI*-RNAi lines exhibited a strong late flowering phenotype under LD^18h^ conditions, demonstrating that all these genes are activators of flowering in LDs in H4 (Figs **1a****, S3A**). However, under SD^12h^ conditions, independent *FvGI-*, *FvCO-*and *FvFT1-*RNAi lines flowered either at the same time or slightly later than H4 (Fig **1b****, S3B**). By contrast, *FvGI*-OE advanced flowering in SD^12h^ but not in LD^18h^ (Figs **1a****, b, S3A, B**).

**Fig. 1.**
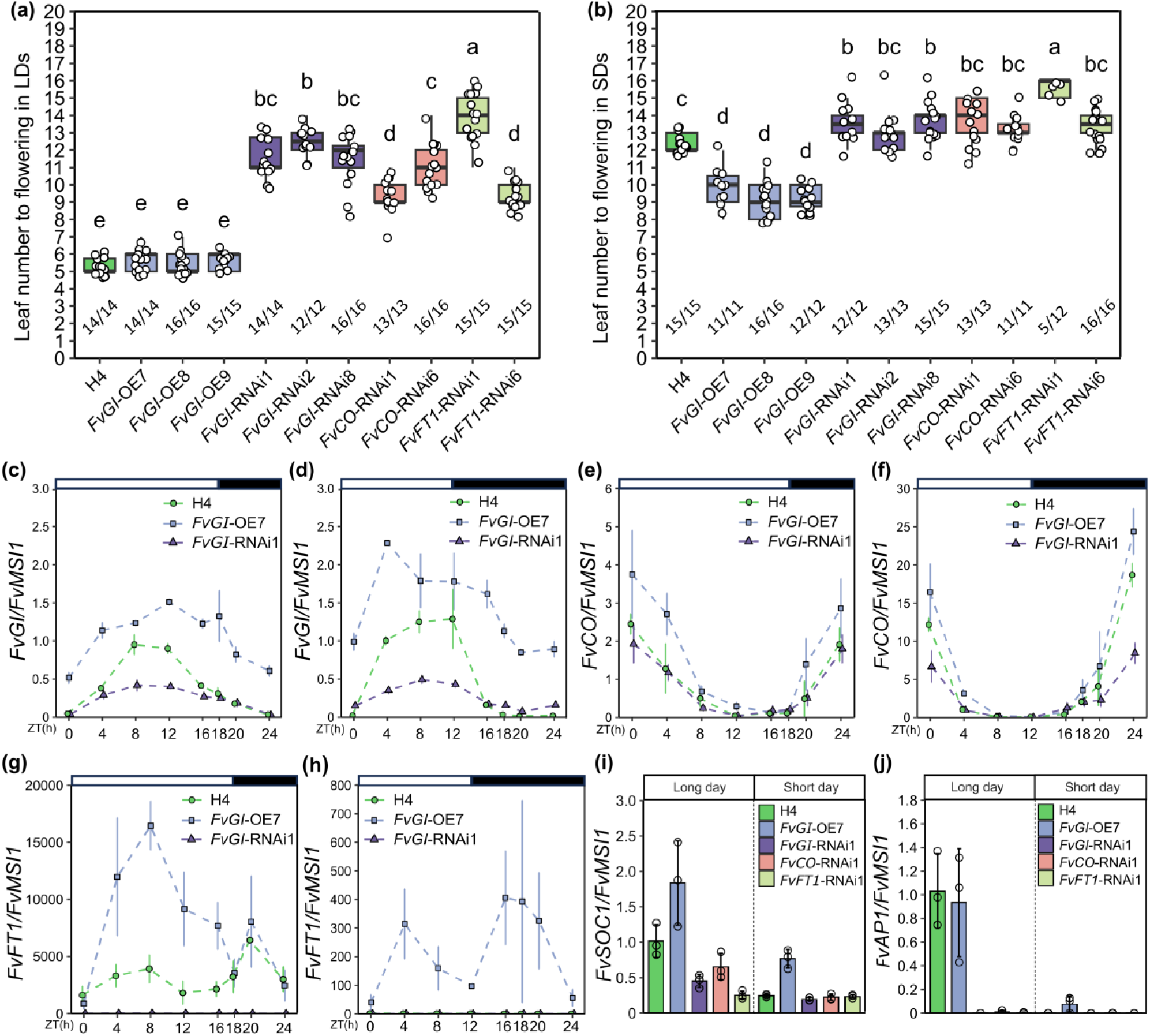
*FvGI*, *FvCO*, and *FvFT1* promote flowering in perpetual flowering *Fragaria vesca*. (a, b) Number of leaves before terminal inflorescence in H4 and indicated transgenic lines under LD^18h^ (a) and SD^12h^ (b) conditions. Numbers under the boxplot indicate the number of flowered plants per the total number of plants in each group. Different letters indicate statistically significant differences as determined by one-way ANOVA followed by Tukey’s HSD multiple comparisons test (P < 0.05). (c-h) Relative expression (mean ± SD, n = 3) of *FvGI*, *FvCO* and *FvFT1* over a 24-h cycle in H4, and *FvGI-*OE and RNAi lines after three weeks under LD^18h^ (c, e, g) or SD^12h^ (d, f, h) conditions. White and black bars above the panels represent light and dark periods, respectively. (i, j) Relative expression (mean ± SD, n = 3) of *FvSOC1* (i) and *FvAP1* (j) in shoot apices of H4, *FvGI-*OE and *FvGI*, *FvCO*, and *FvFT1* RNAi silencing lines. Each biological replicate contains a single shoot apex harvested after 3 weeks under LD^18h^ or SD^12h^ treatment.

Consistent with flowering phenotypes, *FvFT1* expression level was very low in the leaves of *FvGI*, *FvCO* or *FvFT1* RNAi lines sampled four hours after dawn (ZT4) in LD^18h^ (Fig. **S2**). In the *FvGI*-OE lines, *FvCO* and *FvFT1* mRNA levels varied, showing either slight to moderate increase or no difference at ZT4 compared with H4. We next analysed the diurnal expression patterns of *FvGI*, *FvCO* and *FvFT1* in H4 and selected transgenic lines. Under LD^18h^ conditions, *FvGI* expression peaked at ZT8-12 followed by a gradual decline towards the end of the diurnal cycle, while a higher and earlier expression peak (ZT4) was found under SD^12h^ treatment, with a rapid decline towards early night (Fig. **1c, d**). In *FvGI*-OE and RNAi lines, *FvGI* expression remained rhythmic with higher and lower amplitude compared with H4, respectively, under both conditions (Fig. **1c, d**). Furthermore, RNAi silencing of *FvCO* or *FvFT1* did not affect *FvGI* expression compared to H4 indicating that they function downstream of FvGI (Fig. **S4A, B**). *FvCO* expression peaked at dawn/early morning under both conditions in all lines with higher peaks under SD^12h^ treatment. Its expression was elevated in *FvGI*-OE lines, while *FvGI*-RNAi lines showed reduced *FvCO* expression levels in some time points especially in SD^12h^ (Figs **1e****, f, S4C, D)**. In correlation with the observed flowering phenotypes, *FvFT1* mRNA was detected only under LD^18h^ conditions with a minor morning peak (ZT4-ZT8) and a higher evening peak (ZT20) in H4, whereas in *FvGI*-, *FvCO-* and *FvFT1-* RNAi lines, *FvFT1* expression remained nearly undetectable under both photoperiods (Fig. **1g, h, S4E, F**). *FvGI*-OE plants, in contrast, exhibited highly elevated daytime *FvFT1* mRNA levels under LD^18h^ conditions and rhythmic *FvFT1* expression with lower level also under SD^12h^ conditions. The expression of *FvFT2* (FvH4_4g30710), that was previously proposed to function as a SD-activated florigen in *F. vesca* (Gaston *et al*., 2021), was undetectable in all samples (Fig. **S4G**).

Next, we tested the expression of *FvSOC1* (FvH4_7g12700) and the floral identity gene *FvAP1* (FvH4_4g29600) that were previously shown to be regulated by FvFT1 in the shoot apex (Mouhu *et al*., 2013). Under LD^18h^ conditions, *FvSOC1* mRNA level was slightly elevated in *FvGI*-OE7 line and reduced in *FvGI*, *FvCO* and *FvFT1*-RNAi lines compared to H4 (Fig.**1i**). Consistent with late flowering, also *FvAP1* expression was reduced in all RNAi lines (Fig. **1j**). Under SD^12h^ conditions, both *FvSOC1* and *FvAP1* were activated in the early flowering *FvGI*-OE7 line, while low expression levels were found in H4 and RNAi lines. Taken together, our data suggests that FvGI activates the expression of *FvCO* and *FvFT1* in the leaves resulting in early floral transition in LDs in H4.

### *FvGI* represses flowering in seasonal flowering *F. vesca*

In contrast to the perpetual flowering H4, seasonal flowering *F. vesca* contains a functional FvTFL1 (FvH4_6g18480; Koskela *et al*., 2012). We next generated *FvGI*-OE lines in the seasonal flowering accessions, including the reference accession FIN56 and the accession NOR8 that exhibited low *FvFT1* expression and early flowering (Figs **S5A, B, S6A, B**). All plants remained vegetative under non-inductive LD^18h^ conditions. After flowering-inductive SD^12h^ treatment, all NOR8 and FIN56 wild type plants flowered, whereas most of the *FvGI*-OE plants remained vegetative (Fig. **S6C-F**), indicating that FvGI represses flowering in seasonal flowering *F. vesca*. Gene expression analyses confirmed that *FvGI*-OE strongly activated *FvFT1* expression in all lines compared with respective wild type lines, while the effect on *FvCO* mRNA level was milder (Fig. **S7A-G**). In the shoot apex, *FvGI*-OE lines exhibited higher *FvSOC1* and *FvTFL1* expression levels compared to NOR8 (Fig. **S7H-J**), indicating that FvGI may repress flowering by activating *FvTFL1* through *FvSOC1* (Mouhu *et al*., 2013).

To examine the repressor function of *FvGI* in more detail, we subjected wild type NOR8 and two late flowering *FvGI*-OE lines to three photoperiods including LD^16h^, SD^12h^, and SD^8h^ for 5 weeks, followed by flowering observations in LD^18h^. None of the plants flowered after LD^16h^ (Fig. **2a, b**). In SDs, overexpression of *FvGI* shortened the critical photoperiod for flower induction: The SD^8h^ resulted in flowering of 100, 80 and 60% of the NOR8, *FvGI-*OE9 and *FvGI*-OE11 plants, respectively, while after SD^12h^, all NOR8 plants flowered, *FvGI*-OE9 remained vegetative and 20% of the *FvGI*-OE11 plants flowered. *FvGI*-OE plants flowered about one week later and with two more leaves than NOR8 after the SD^8h^ treatment. Slower floral transition in *FvGI*-OE plants was also visible in the meristems. The SEM images, taken at the end of the treatments, showed more developed primary flowers in NOR8 compared with transgenic lines (Fig. **S8**).

**Fig. 2.**
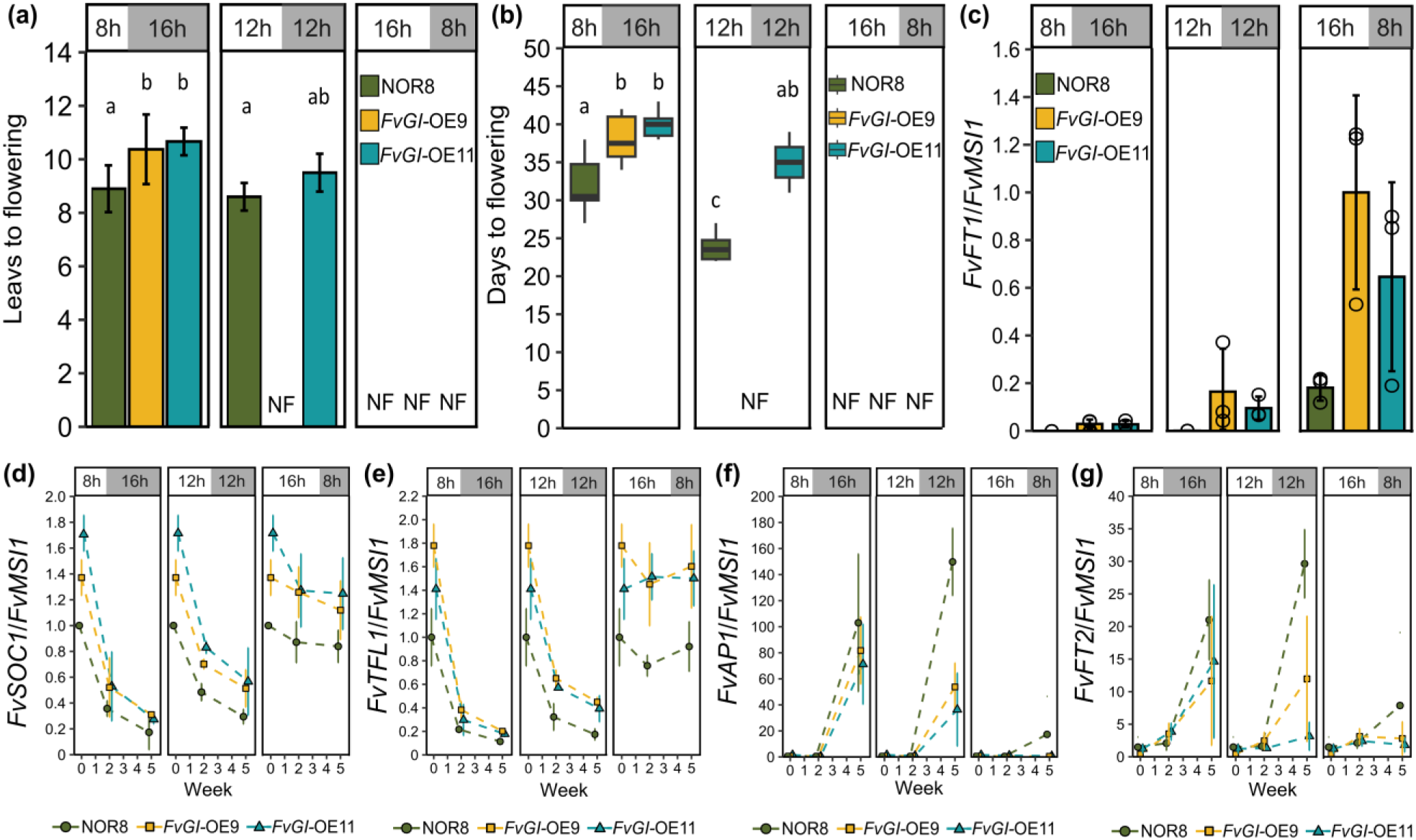
*FvGI* represses flowering in seasonal flowering *Fragaria vesca*. (a) Number of leaves before terminal inflorescence in NOR8 and *FvGI-OE* lines under indicated conditions. (b) Flowering time of NOR8 and *FvGI*-OE lines indicated as days to anthesis from the end of indicated treatments. In A and B, values indicate mean ± SD (*n* = 10); NF= no flower. Different letters indicate statistically significant differences as determined by one-way ANOVA followed by Tukey’s HSD multiple comparisons test (P < 0.05). Clonally propagated plants were pre-cultured under LD^18h^ conditions for four weeks, after which the plants were subjected to SD^8h^, SD^12h^ or LD^16h^ at 17 °C for five weeks followed by phenotyping under LD^18h^. (c) *FvFT1* expression (mean ± SD, n = 3-4) in leaf samples collected at ZT4 after five weeks under the indicated photoperiods. (d-g) *FvSOC1* (d), *FvTFL1* (e), *FvAP1* (f) and *FvFT2* (g) expression (mean ± SD, n = 3) in shoot apices after 0, 2 and 5 weeks under indicated photoperiods. Three shoot apices were harvested for each biological replicate.

Gene expression analysis showed that *FvFT1* expression increased in leaves of NOR8 and *FvGI*-OE lines towards longer daylengths being higher in *FvGI*-OE lines than in the wild type in all photoperiods at ZT4 (Fig. **2c**). Higher *FvFT1* expression was associated with elevated *FvSOC1* and *FvTFL1* expression in the shoot apices (Fig. **2d, e**). Both *FvSOC1* and *FvTFL1* were downregulated under shorter photoperiods with the SD^8h^ causing stronger repression than the SD^12h^. The strong downregulation of *FvSOC1* and *FvTFL1* was observed also in *FvGI*-OE lines, but their expression levels remained always higher than in the wild type. Consistent with flowering results, the floral meristem identity gene *FvAP1* was strongly upregulated in the end of the treatments in all genotypes under SD^8h^ and in NOR8 under SD^12h^ (Fig. **2f**). The expression pattern of *FvFT2* closely followed the expression of *FvAP1* in the shoot apex, but its expression was not detected in leaves of NOR8 or its *FvGI*-OE lines (Figs **2g****, S9A, B**), suggesting that FvFT2 may function in early floral development. Taken together, our results indicate that, in seasonal flowering *F. vesca*, *FvGI* activates the expression of *FvFT1* in the leaves and *FvSOC1* and *FvTFL1* in the shoot apex, repressing floral transition in LDs.

### Genetic interactions in the regulation of photoperiodic flowering

Gene expression analyses in transgenic lines suggested that FvGI functions upstream of *FvCO* and *FvFT1* (Figs **2e-h**, **3c, S7D-G**). To test the genetic interactions of these genes, we crossed NOR8 *FvGI*-OE line#9 and wild type NOR8 (control) with H4, and *FvCO-* and *FvFT1*-RNAi lines in H4 background. Half of the NOR8 *FvGI*-OE × H4 F_1_ hybrids remained vegetative after the SD^12h^ treatment, and the rest of the plants flowered later than NOR8 × H4 controls (Figs **3a****, b, S10**), confirming that *FvGI*-OE represses flowering in the hybrids. However, the silencing of *FvCO* or *FvFT1* resulted in equal early flowering in both NOR8 or NOR8 *FvGI*-OE hybrids, demonstrating that both *FvCO* and *FvFT1* are required for the late flowering of NOR8 *FvGI*-OE plants. While NOR *FvGI*-OE × H4 hybrids showed elevated *FvFT1* expression levels compared with the NOR8 × H4 hybrids, silencing of *FvCO* rendered *FvFT1* undetectable regardless of *FvGI*-OE, indicating that the activation of *FvFT1* by FvGI requires *FvCO*. Furthermore, crossing of NOR8 or NOR8 *FvGI*-OE line with H4 *FvFT1*-RNAi line did not affect *FvCO* expression. These results support the presence of *FvGI*-*FvCO*-*FvFT1* photoperiodic flowering pathway in *F. vesca*, where *FvCO* and *FvFT1* are fully epistatic to *FvGI*.

**Fig. 3.**
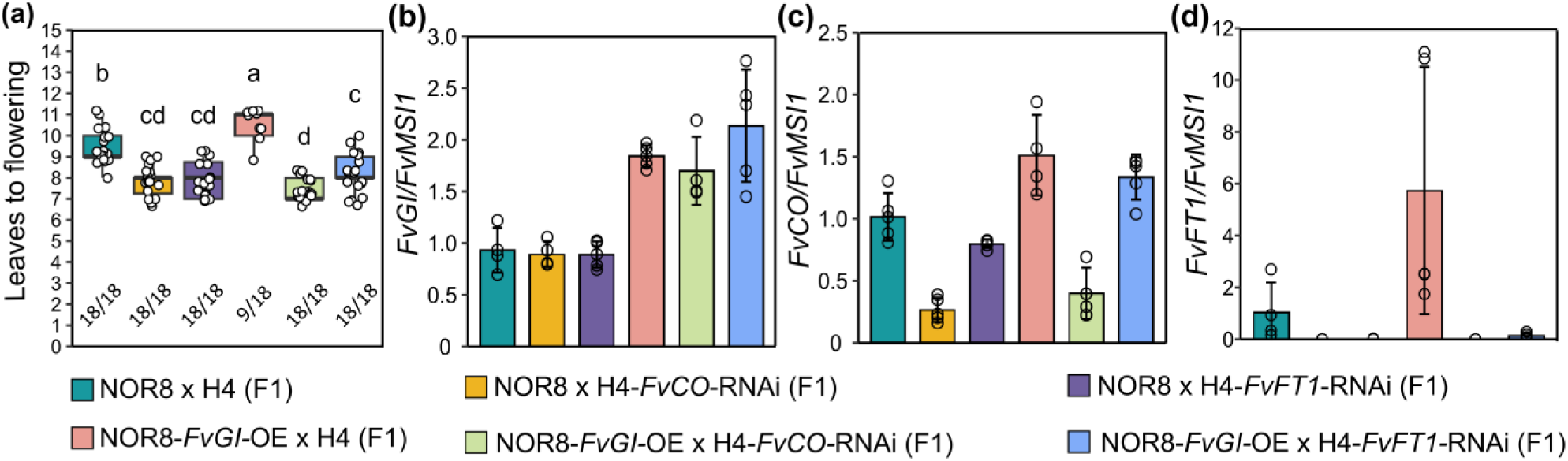
The *FvGI–FvCO–FvFT1* pathway regulates flowering time in seasonal flowering *Fragaria vesca*. (a) Flowering time of indicated F_1_ hybrids shown as the number of leaves before the terminal inflorescence. Four-week-old plants, clonally propagated from hybrid seedlings were subjected to SD^12h^ at 17°C for eight weeks followed by phenotyping under LD^18h^. Numbers under the boxplot indicate the number of flowered plants per the total number of plants in each group. Different letters indicate statistically significant differences as determined by one-way ANOVA followed by Tukey’s HSD multiple comparisons test (P < 0.05). (b-d) Relative expression of *FvGI* (b), *FvCO* (c) and *FvFT1* (d) in leaves of F_1_ lines. Leaf samples containing one leaflet per biological replicate were harvested at ZT4 under LD^18h^ conditions. Data are mean ± SD (n = 4-5).

### *FvTFL1* reverses the effect of *FvFT1* on flowering time

Because *FvGI*-OE was found to activate *FvFT1* in the leaves and *FvTFL1* in the shoot apex of seasonal flowering *F. vesca* (Fig. **2c, e**), we tested if FvFT1 activated *FvTFL1*. To this end, we crossed seasonal flowering accessions, the reference accession FIN56 and Italian accession IT17 with naturally high *FvFT1* expression level (Fig. **S5**), with H4 and H4 *FvFT1*-RNAi line and analysed the F_1_ hybrids. *FvFT1*-RNAi hybrids with strongly reduced *FvFT1* mRNA levels flowered early after SD^12h^ treatment, and 40% of IT17 × H4 *FvFT1*-RNAi and 67% of FIN56 × H4 *FvFT1*-RNAi plants flowered also under non-inductive LD^18h^ conditions (Fig. **4a, b**). The expression levels of *FvSOC1* and *FvTFL1* were lower in the shoot apices of *FvFT1*-RNAi hybrids than in the control plants in LD^18h^ conditions (Figs **4c****, S11**). After three weeks of SD^12h^ treatment, *FvTFL1* expression decreased in all plants but remained higher in crosses with the wild type H4 compared with H4 *FvFT1*-RNAi hybrids. At the same time, *FvAP1* exhibited high expression levels only in the *FvFT1*-RNAi hybrids, confirming that flower induction had happened in these plants but not in the wild type hybrids (Fig. **4d**). Taken together, our results suggest that *FvFT1* activates *FvTFL1* expression in the shoot apex to repress flowering in seasonal flowering *F. vesca*, and plants with higher *FvFT1* and *FvTFL1* mRNA levels require longer inductive treatment to reduce the expression level of *FvTFL1* below a critical threshold for floral induction.

**Fig 4.**
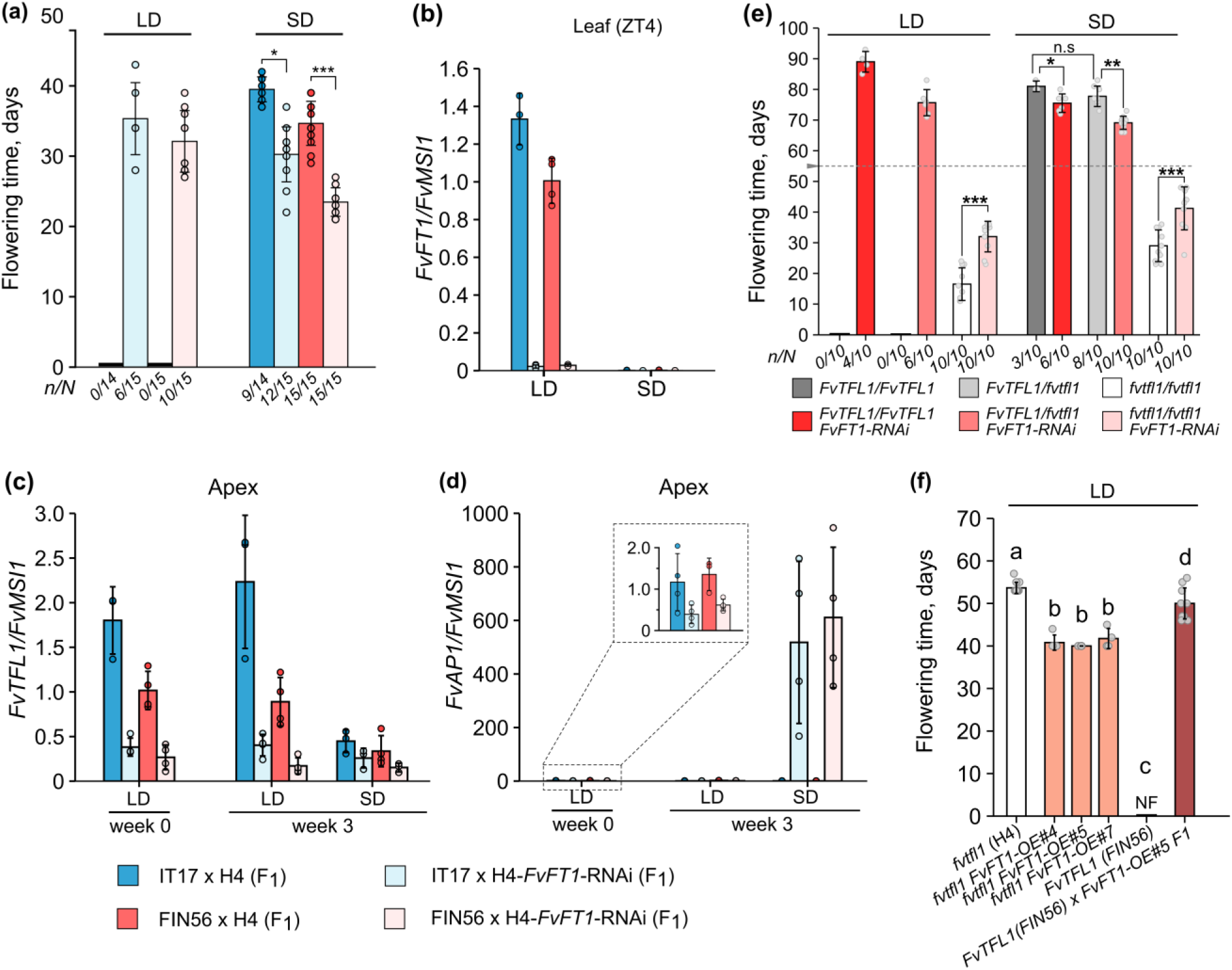
*FvFT1/FvTFL1* epistasis controls flowering in *Fragaria vesca*. (a) Flowering time of F_1_ hybrids between the seasonal flowering genotype IT17 or FIN56 with functional *FvTFL1* alleles and the perpetual flowering genotype H4 (*fvtfl1*) carrying *FvFT1*-RNAi construct after six weeks under SD^12h^ conditions. F_1_ crosses with non-transgenic H4 plants were used as controls. Asterisks indicate significant differences between groups at selected time points (***, p < 0.001; **, p < 0.01; *, p < 0.05; t-test). (b-d) Expression of *FvFT1* (b) in leaf samples and *FvTFL1* (c) and *FvAP1* (d) in shoot apices under LD^18h^ conditions and after three weeks of indicated treatments at ZT4 time point. Insert panel in D shows a close view of initial (before the start of treatment) expression levels of *FvAP1* at week 0. (e) Flowering time of IT17 × *FvFT1*-RNAi F_2_ plants segregating for *FvFT1-*RNAi construct, and *FvTFL1* and *fvtfl1* alleles after eight weeks under SD^12h^ conditions. y-axis shows days since the beginning of the treatments; dashed grey line marks the end of the indicated treatments after which all the plants were moved to LD^18h^ conditions. Numbers under x-axis denote the number of flowered plants (*n*) and total number of plants (*N*) in each group. (f) Flowering time of H4 and FIN56 wild type plants and their *FvFT1-*OE lines under LD^18h^ conditions. In e and f, data are mean ± SD (n = 10). Asterisks (e) or different letters (f) indicate the significant differences between selected groups (***, p < 0.001; **, p < 0.01; *, p < 0.05; Tukey’s HSD test).

To obtain direct evidence of the interaction of the photoperiodic pathway with *FvTFL1* we segregated *FvTFL1* alleles and *FvFT1-*RNAi insertion in IT17 × H4 *FvFT1*-RNAi F_2_ lines and observed the flowering time. Under both LD^18h^ and SD^12h^ conditions, *fvtfl1* homozygous mutants with the *FvFT1*-RNAi construct flowered later than the *fvtfl1* mutants without the *FvFT1-*RNAi construct, confirming that FvFT1 activates flowering in the absence of functional FvTFL1 (Fig. **4e**). However, the presence of one or two alleles encoding functional FvTFL1 resulted in SD-dependent flowering and reverted the function of FvFT1; the plants with the *FvFT1*-RNAi construct flowered earlier than the controls. In addition, approximately half of the *FvFT1*-silenced plants homozygous or heterozygous for functional FvTFL1 flowered even under non-inductive LD conditions. A higher proportion of flowering plants and earlier flowering of *FvTFL1/fvtfl1* heterozygotes compared with plants containing two wild type alleles, supported the dosage-dependent effect of *FvTFL1* on flowering in *F. vesca* (Fig. **4e**).

Altogether, our results indicate that flowering regulation in *F. vesca* involves an epistatic interaction between the *FvGI-FvCO-FvFT1* photoperiodic pathway and *FvTFL1*. When the functional *FvTFL1* allele is present, FvFT1 prevents flowering in LDs by promoting *FvTFL1* expression, while in the absence of functional *FvTFL1*, FvFT1 functions as a florigen that activates the expression of *FvAP1* thus promoting flowering. Even in plants with functional FvTFL1, FvFT1 retains florigenic activity, as its RNAi silencing slightly reduces *FvAP1* expression (Fig. **4d**), but this activity is masked by the *FvTFL1* activation. Moreover, the expression of *FvFT1* specifically in the leaves (Koskela *et al*., 2012) is critical for flowering repression mechanism presented here, because ectopic expression of *FvFT1* by the 35S promoter induces flowering even in the presence of *FvTFL1* (Fig. **4f**).

### *FvFT1* contributes to natural variation in flowering time

Finally, we explored natural flowering time variation in a European collection of *F. vesca* accessions that were recently shown to form separate eastern and western genetic clusters occupying different climatic niches (Toivainen *et al*., 2026). After flower induction under SD^12h^ in a greenhouse or field conditions in autumn in Helsinki, a wide variation in flowering time was observed with the eastern accessions flowering significantly earlier than the western ones suggesting adaptive variation (Figs **5a-c**).

**Fig. 5.**
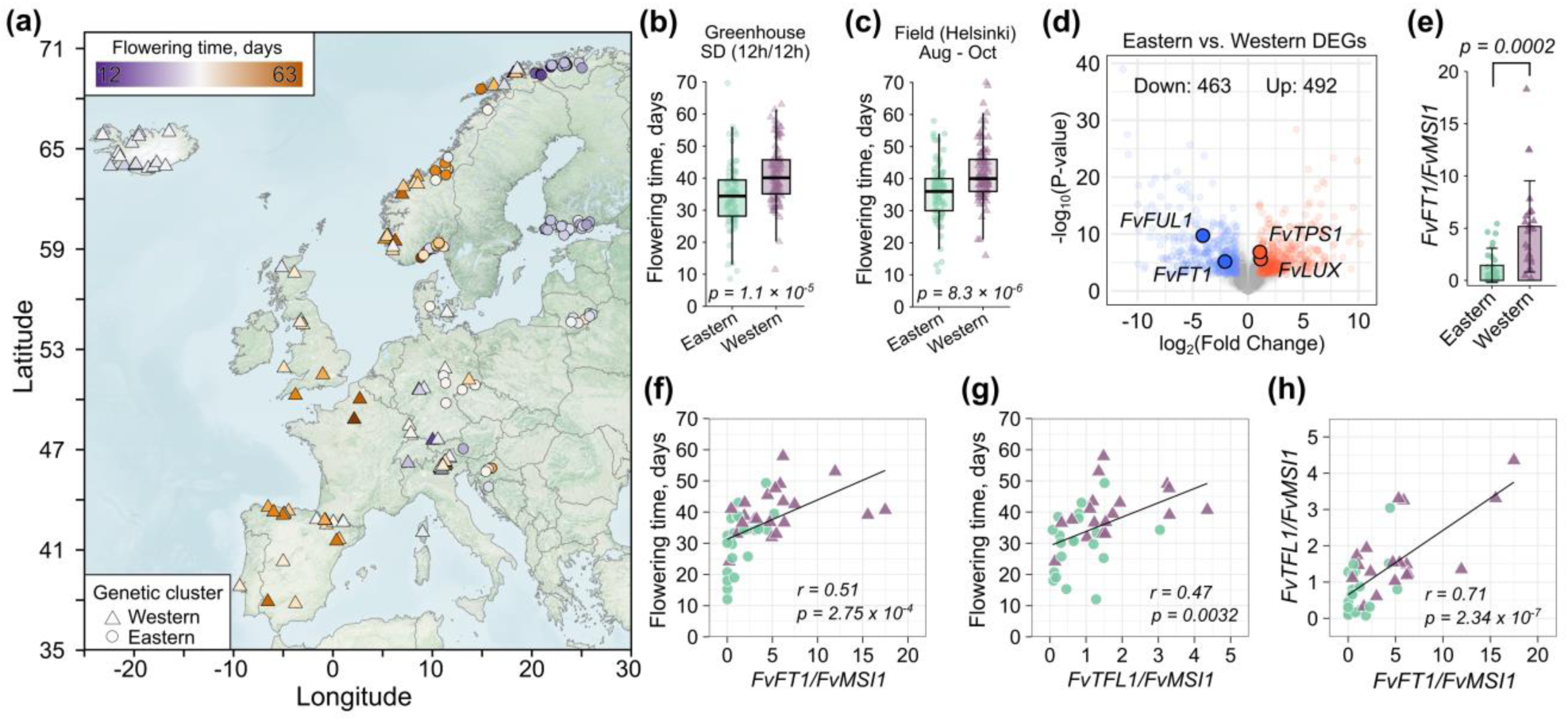
Natural variation of flowering time correlates with *FvFT1* and *FvTFL1* expression in European *F. vesca* accessions. (a) Map showing collection sites of woodland strawberry accessions and their flowering time and genetic clusters (Toivainen *et al*. 2026). (b-c) Differences in flowering time between the accessions of eastern and western genetic clusters after flower induction under SD^12h^ conditions in a greenhouse (b; 179 accessions) or in field (c; 181 accessions). (d) Differentially expressed genes (DEGs) in leaves (LD^18h^, ZT4) of six western and six eastern accessions. (e) Mean expression levels of *FvFT1* in 25 eastern and 23 western accessions. (f-g) Pearson’s correlation between flowering time and *FvFT1* (f) or *FvTFL1* (g) expression levels using the expression data shown in e and h. (h) Pearson’s correlation between the relative expression of *FvFT1* in leaves and *FvTFL1* in shoot apices (LD^18h^, ZT16) in 21 eastern and 19 western accessions.

RNA sequencing from leaves of six eastern and six western accessions identified 955 genes differentially expressed between the genetic clusters (Dataset **S1**). Among these, four candidate flowering time genes were found including *FvTPS1* (*TREHALOSE-6-PHOSPHATE SYNTHASE1*, FvH4_3g06590)*, FvLUX* (*LUX ARRHYTHMO,* FvH4_3g36890), *FvFUL1* (*FRUITFUL*, FvH4_5g13500) and *FvFT1* (Figs **5d****, S12A-D**). Both *LUX* and *TPS1* are known to affect *FT* expression in Arabidopsis (Hazen et al., 2005; Wahl et al., 2013) and a FUL homolog in barley (Sasani et al., 2009), while their functions in *F. vesca* remain to be clarified. The expression of *FvFT1* was typically lower in the accessions of the earlier flowering eastern cluster (Fig. **S12A**). We verified this by RT-qPCR analysis in a larger set of accessions (25 eastern and 23 western), spanning the geographical range of European *F. vesca* (Figs **5e****, S5**). Furthermore, the expression of *FvFT1* in leaves of accessions strongly correlated with *FvTFL1* expression levels in the shoot apex and the flowering time (Figs **5f-h****, S12G-J**). Although *FvGI* and *FvCO* expression levels did not consistently differ between the two clusters in the RNAseq data (Fig. **S12E, F**), their expressions were correlated among a larger set of accessions, as were *FvCO* and *FvFT1* (Fig. **S12K, L**). Altogether, these findings indicate that the *FvGI-FvCO-FvFT1-FvTFL1* photoperiodic pathway contributes to flowering time variation across geographic and climatic ranges in *F. vesca* accessions, with strong correlations between flowering time and latitude or winter temperature of plant collection sites (Fig. **S12M**).

## DISCUSSION

Here, we studied photoperiodic flowering in *F. vesca*, a perennial species with a wide geographical distribution in the northern hemisphere (Hilmarsson *et al*., 2017; Toivainen *et al*., 2026). Previous studies showed that the floral repressor FvTFL1 confers the SD requirement of flower induction in *F. vesca*, while the *fvtfl1* mutant flowers early in LDs (Koskela *et al*., 2012). We show that a conserved *FvGI-FvCO-FvFT1* pathway expressed in the leaves activates LD-flowering in the *fvtfl1* mutant, while in seasonal flowering, wild type plants this pathway is rewired to activate *FvTFL1* in the shoot apex, reversing the photoperiodic response. Due to this connection, wild type plants require SDs for flower induction that occurs after the downregulation of *FvFT1* and *FvTFL1* in autumn. In natural accessions, *FvFT1* expression correlates positively with *FvTFL1* mRNA levels and is associated with differences in flowering time in plants collected across Europe, suggesting that a flower-repressive photoperiodic pathway contributes to flowering time adaptation across geographic and climatic ranges.

### Epistatic interaction between *FvFT1* and *FvTFL1* reverses photoperiodic response

Homologs of FT have been shown to induce flowering in both LD and SD plants (Kobayashi *et al*., 1999; Faure *et al*., 2007; Tamaki e*t al.*, 2007; Komiya *et al*., 2009; Kong *et al*., 2010), while TFL1 represses flowering (Bradley *et al*., 1997; Wang *et al*., 2011), possibly thorough competing for binding with FD transcription factor that is needed for the regulation of their target genes (Zhu *et al*., 2020). Multiple lines of evidence suggested different mechanisms in the regulation of photoperiodic flowering in *F. vesca: FvFT1* is expressed in leaves in LDs when *FvTFL1* is also activated in the shoot apex to repress flowering, and floral transition occurs in SDs when *FvTFL1* is downregulated and *FvFT1* expression is undetectable (Koskela *et al*., 2012). In *fvtfl1* mutant, however, *FvFT1* promotes flowering in LDs and is fully responsible for this photoperiodic response (Koskela *et al*., 2012; Rantanen *et al*., 2014; Kurokura e*t al.*, 2017). By segregating *FvTFL1* alleles and the *FvFT1*-RNAi insertion in an F_2_ crossing population we demonstrated that *FvTFL1* is epistatic to *FvFT1* in the photoperiodic flowering. All F_2_ lines homozygous for *fvtfl1* alleles flowered early in LDs, with RNAi-silencing of *FvFT1* delaying flowering indicating that FvFT1 promotes flowering in the *fvtfl1* mutant. In contrast, when one or two functional *FvTFL1* alleles were present, flowering became SD-dependent, and the effect of *FvFT1* was reversed to repressor of flowering. *FvFT1*-RNAi plants flowered earlier than controls after the SD treatment, and a substantial proportion of *FvFT1*-RNAi plants flowered even in non-inductive LDs.

To understand the mechanism of the epistatic interaction between *FvFT1* and *FvTFL1*, we studied gene expression in *FvFT1*-RNAi lines and natural accessions of *F. vesca*. We found lower expression level of *FvTFL1* in *FvFT1*-RNAi lines compared with wild type plants and positive correlation between *FvFT1* and *FvTFL1* mRNA levels in natural accessions, suggesting that FvFT1 activates *FvTFL1* in the SAM under LDs to repress flowering in *F. vesca*. The activation of *FvTFL1* occurs at least partially through *FvSOC1* that is regulated by FvFT1 (Mouhu *et al*., 2013) (Figs **1i****, S7G, H)**. Recent work in Arabidopsis showed that FT-activated LEAFY (LFY) upregulated *TFL1* to maintain indeterminate inflorescence meristem (Huang *et al*., 2026). Further analysis of four *LFY* genes present in *F. vesca* genome is needed for testing if LFY has similar role in *F. vesca* (Zhang *et al*., 2023). The *FvFT1/FvTFL1* module has a quantitative effect on floral transition and flowering time, because *FvFT1*-RNAi lines showed early upregulation of *FvAP1* and early flowering, and *FvTFL1/fvtfl1* heterozygote plants flowered earlier than *FvTFL1* homozygotes (Samad *et al*., 2017, Fig. **4e**). Overall, the level of *FvFT1/FvTFL1* expression is likely to drive the strength of floral repression in LDs to control the duration of inductive SD period needed for floral transition. However, the mechanism of flower induction in SDs remains elusive. Previous study found the expression of *FvFT2* in leaves and suggested that FvFT2 could induce flowering in SDs (Gaston *et al*., 2021). We could not detect *FvFT2* expression in the leaves of tested accessions and transgenic lines under any conditions, but *FvFT2* was upregulated in the shoot apex in parallel with *FvAP1* indicating that it plays a role in early floral development instead of floral induction. Possible role of SQUAMOSA PROMOTER-BINDING PROTEIN-LIKE (SPL) transcription factors, that induce flowering in SDs in *A. thaliana* and *A. alpina* (Roggen *et al*., 2025), should be tested in *F. vesca*.

### *FvGI-FvCO-FvFT1* pathway controls photoperiodic flowering

*CO/FT* module has been shown to control photoperiodic flowering in annual models Arabidopsis and rice (Valverde *et al*., 2004; Ishikawa *et al*., 2011), and among perennials, in perpetual flowering mutants of *F. vesca* and *A. alpina* (*fvtfl1* and *pep1* mutants, respectively; Kurokura *et al*., 2017; Sashidhar & Coupland, 2026). Here, we extended these studies to *FvGI* and demonstrated that all three genes are required for the photoperiodic flowering of *F. vesca*. In RNAi lines of *FvGI, FvCO* and *FvFT1* in H4 background (*fvtfl1* mutant; Koskela *et al*., 2012), *FvFT1* expression levels remained below/close to the detection limit in LDs, and the lines almost completely lost the early LD flowering observed in H4. *FvGI*-OE, in contrast, increased *FvFT1* expression resulting in early flowering specifically in SDs in H4 background and delayed SD-flowering in plants possessing functional *FvTFL1* (NOR8, FIN56), consistent with the epistatic effect of *FvTFL1* to *FvFT1*. Furthermore, late flowering phenotype of NOR8 *FvGI*-OE plants was completely reversed by silencing of *FvCO* or *FvFT1* that also reduced *FvFT1* expression close to the detection limit, demonstrating that *FvCO* and *FvFT1* were epistatic to *FvGI*. Together with previous research on *FvCO* function (Kurokura *et al*., 2017), these findings established the *FvGI-FvCO-FvFT1* pathway with a major role in mediating the LD signal to activate flowering in *F. vesca fvtfl1* mutants. In wild type plants, this pathway activates *FvTFL1*, and SDs are required to suppress the whole pathway to induce flowering. In other SD plants, different mechanisms repress flowering in LDs. In rice, Ghd7 blocks *Hd1* (CO-homolog) function and in soybean E1 and GmCOL1a/b repress *FT2a/5a* and flowering (Xia *et al*., 2012; Cao *et al*., 2015; Xu *et al*., 2015; Zong *et al*., 2021).

Sequences and diurnal expression rhythms of *GI* homologs are largely conserved in both LD and SD plants, suggesting broadly conserved roles in photoperiodic signalling (Fowler et al., 1999; Hayama et al., 2003; Hecht et al., 2007; Li et al., 2013; Deng et al., 2015, **Fig. S1**). In Arabidopsis, GI promotes *CO* transcription in LDs, allowing CO protein to peak in the evening to activate *FT* (Valverde *et al*., 2004; Sawa *et al*., 2007). However, *FvCO* mRNA peaks at dawn with a broader expression phase in LDs, the pattern resembling that of Arabidopsis *COL1/COL2*, pea *PsCOLa*, and soybean *GmCOL1a/b* (Ledger *et al*., 2001; Hecht *et al*., 2007; Wu *et al*., 2014). This *FvCO* expression pattern is needed for normal *FvFT1* expression peaks in the morning and late evening, because both peaks are lost in *FvCO* RNAi lines (Kurokura *et al*., 2017). Similarly, *FvFT1* expression peaks were lost in *FvGI*-RNAi lines, although *FvCO* mRNA levels were not clearly affected, indicating that FvGI regulates *FvFT1* expression mainly through posttranscriptional regulation of FvCO, which is further supported by the observations that both *FvGI*-OE and *FvCO*-OE had only a weak effect on *FvFT1* mRNA levels in SDs, compared with LDs (Kurokura *at el.*, 2017, Fig. **1e-h**). Such a mechanism is consistent with a LD-dependent stabilization of CO in Arabidopsis that involves FKF1 and GI (Valverde *et al*., 2004; Song *et al*., 2012; 2014, Hwang, *et al*., 2019). In conclusion, we show strong evidence for the *FvGI-FvCO-FvFT1* photoperiodic pathway in *F. vesca*. Further studies are needed to understand posttranscriptional regulation of FvCO and a possible role of *FvFKF1*, that has overlapping expression phase with *FvGI* (Kurokura *et al*., 2017), in photoperiodic flowering of *F. vesca*.

### Flowering time variation in European *F. vesca*

*F. vesca* occupies a wide geographical range from Mediterranean to arctic region in the northern hemisphere (Hilmarsson *et al*., 2017). Our data indicates that flowering time variation in *F. vesca* accessions collected across Europe is associated with their population structure, which consists of two major genetic clusters. These western and eastern clusters originate from different glacial refugia and inhabit distinct climates across temperature seasonality gradient (Toivainen *et al*., 2026). We found that eastern accessions flowered significantly earlier than western accessions under both SD and field conditions, with flowering time most strongly correlating with mean annual and winter temperatures of plant collection sites. Also, the latitudinal correlation was strong, with accessions from arctic regions, particularly in the east, showing extreme early flowering. Such pattern of flowering time variation can be attributed to the length of the growing season that creates strong selection pressure on reproductive traits (Austen *et al*., 2017), suggesting adaptive divergence in flowering regulation.

Reduced activity of the photoperiodic pathway was suggested to contribute to adaptation to short growing seasons in both annual and perennial plants (Hyun *et al*., 2019). Consistent with this idea, we found that decreasing expression levels of *FvFT1* and *FvTFL1* correlated with earlier flowering in European *F. vesca* accessions. While *FvFT1* and *FvTFL1* mRNA levels showed a strong positive correlation with clear association with genetic clusters, weaker positive *FvGI/FvCO* and *FvCO/FvFT1* correlations were found, and there was no clear east-west pattern in *FvGI* or *FvCO* expression, suggesting that *FvFT1* expression variation associated with geography and population structure depends on mechanisms other than the regulation of *FvGI* or *FvCO* transcription. However, arctic accessions (NOR14 and NOR15) from highly differentiated Kåfjord population (Toivainen *et al*., 2026), that flowered first in different experiments, exhibited clearly reduced *FvCO* expression and extremely low *FvFT1* mRNA levels, reflecting population-specific responses to photoperiods. Other mechanisms may include cis-regulatory variation of *FT* that has been reported to contribute to flowering time variation in Arabidopsis and soybean (Liu *et al*., 2014, Rosas *et al*., 2014, Bao *et al*., 2019, Chen *et al*., 2020). While FvGI and FvCO play major roles as activators of *FvFT1* expression, studies on additional candidate genes, including *FvTPS1*, *FvLUX*, and *FvFUL1* identified in our RNA sequencing work, are needed for more detailed understanding of natural flowering time variation and underlying regulatory network in *F. vesca*.

Based on the findings of this study, we propose a model on the photoperiodic regulation of flowering in *F. vesca* (Fig. 6). *FvGI* and *FvCO* are needed for the activation of *FvFT1* in leaves in LDs, and they constitute a major photoperiodic response pathway of the species (Fig. **6a**). In perpetual flowering *fvtfl1* mutant, this pathway activates *FvSOC1* and *FvAP1* in the shoot apex, resulting in rapid flowering in LDs. Functional *FvTFL1,* that is present in all natural accessions tested in this work, reverses the photoperiodic response, because the photoperiodic pathway activates *FvTFL1* at least partially through *FvSOC1*, overriding *FvAP1* activation and repressing flowering (Fig. **6a**). Upstream of *FvFT1*, *FvGI* exhibits daytime specific expression pattern with a peak in the afternoon, while *FvCO* mRNA peaks at dawn/early morning, with a longer tail of expression during LDs. As the effect of FvGI on *FvCO* mRNA levels was weak, stabilization of FvCO protein is a potential mechanism for the activation of *FvFT1* in LDs (Fig. **6b**). Finally, natural variation in *FvFT1* expression level, which is positively correlated with *FvTFL1* expression, may contribute to natural flowering time variation associated with the length of the growing season in European *F. vesca* accessions (Fig. **6c**).

**Fig. 6.**
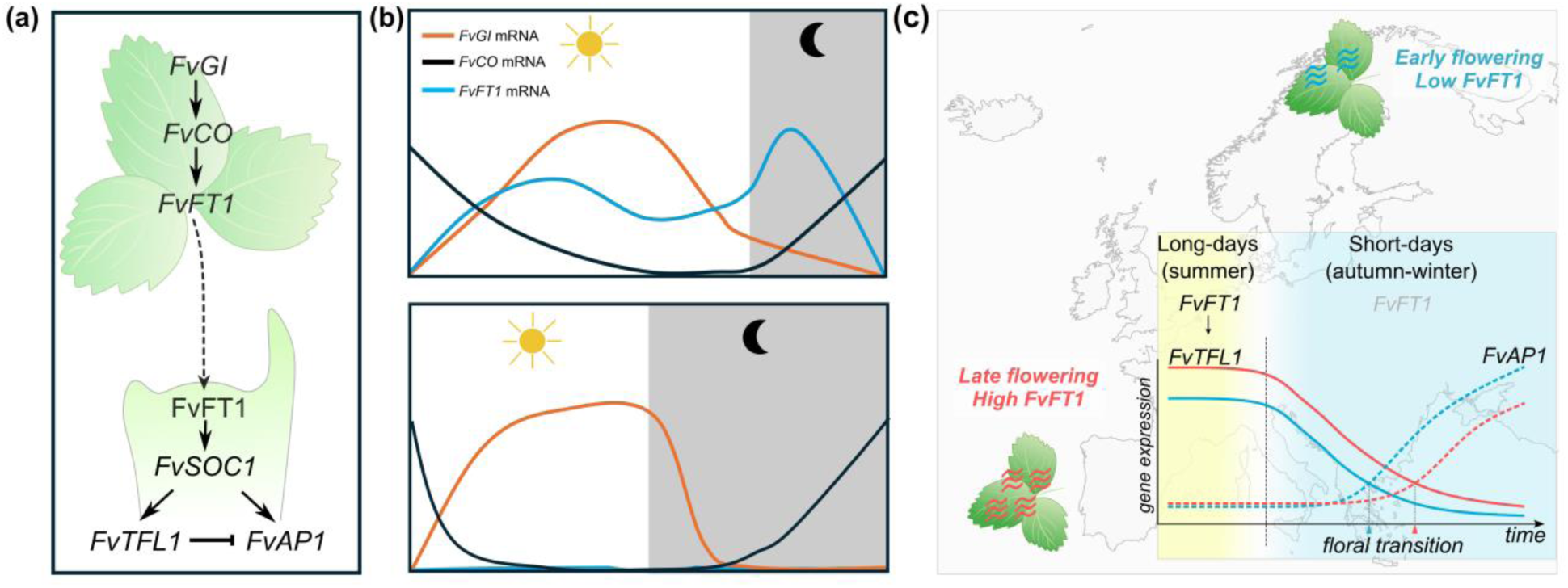
The regulation of photoperiodic flowering in *Fragaria vesca.* (a) A genetic pathway model for photoperiodic flowering in *F. vesca*. Leaf-expressed *FvGI-FvCO-FvFT1* pathway promotes the expression of *FvTFL1* and *FvAP1* in the shoot apical meristem. In the presence, of functional FvTFL1, the activation of the pathway in long days (LD) blocks flowering, while in *fvtfl1* mutant, *FvAP1* is activated resulting in LD-flowering. (b) Expression patterns of leaf-expressed photoperiodic pathway genes. *FvGI* is activated during the day and repressed during night, while *FvCO* peaks at dawn. Longer tail of *FvCO* expression peak is found in LDs, where *FvFT1* exhibits two expression peaks, while in short days (SD), *FvFT1* mRNA is undetectable. Protein level evidence is needed to understand how FvCO creates such *FvFT1* expression pattern. (c) *FvFT1/FvTFL1* module controls natural variation in flowering time. Higher expression level of *FvFT1* is found in the leaves of western/southern compared with eastern/northern accessions. Due to activation of *FvTFL1* by FvFT1, positive correlation is found in their expression levels across accessions. Delayed downregulation of *FvTFL1* (solid lines) in late flowering plants results in delayed floral transition, as indicated by the activation of *FvAP1* (dashed lines), contributing to natural variation in flowering time.

## ACKNOWLEDGEMENTS

This research was funded by the Research Council of Finland (Grants No. 317306 and 368864 to TH). QZ and GF were supported by grants from the China Scholarship Council (QZ, Grant No. 202007960007; GF, Grant No. 201706510014). SL was supported by a personal grant from the Finnish Cultural Foundation (Grant No. 00221168). We are grateful to Eija Takala and Marjo Kilpinen for laboratory assistance, and to Katriina Palm for maintaining the plant materials.

## AUTHOR CONTRIBUTIONS

QZ, SL, EAK and TH designed the research. QZ, SL, TT, TK and GF carried out the experimental work. SL, TT and GF analysed RNA sequencing data. TH, EAK and PE supervised the study. The manuscript was written by QZ and TH with input from all authors.

## DATA AVAILABILITY

All the raw sequence reads from RNA sequencing are available at NCBI under Bio-project ID: ID will be added upon acceptance of the manuscript.

## SUPPORTING INFORMATION

Fig. S1. Phylogenetic analysis and multiple sequence alignment of GI homologs.

Fig. S2. Gene expression in H4 and transgenic lines.

Fig. S3. Flowering time of H4 and transgenic lines.

Fig. S4. Diurnal gene expression rhythms in H4 and transgenic lines.

Fig. S5. Flowering time and *FvFT1* expression in natural accessions of *Fragaria vesca*.

Fig. S6. Gene expression and phenotypes of indicated *Fragaria vesca* genotypes.

Fig. S7. Expression of flowering time genes in NOR8 and *FvGI* overexpression lines.

Fig. S8. Scanning electron microscope images of *Fragaria vesca* meristems.

Fig. S9. Expression of *FvFT1* and *FvFT2* in NOR8 and *FvGI* overexpression lines.

Fig. S10. Flowering time of indicated hybrids.

Fig. S11. Expression of flowering genes in indicated hybrids.

Fig. S12. Comparison of gene expression levels in *Fragaria vesca* accessions and correlations between gene expression, flowering time, and bioclimatic and geographical variables.

Table S1. Hybrid plants used in this study.

Table S2. Woodland strawberry accessions used in this study.

Table S3. Primers used in this study.

Table S4. Protein sequences used for the phylogenetic analysis of GI.

Dataset S1. RNA sequencing data.

