## Supporting Information Figures S1-12 for "*FvTFL1* reverses the function of *FvGI-FvCO-FvFT1* pathway in the photoperiodic flowering of woodland strawberry"

A

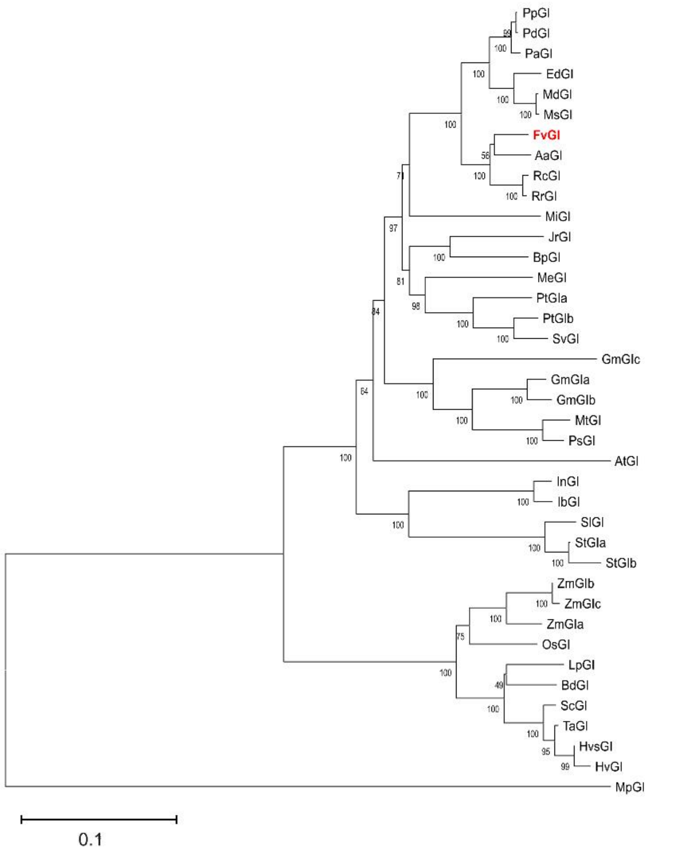
PpGI
PdGI
PaGI
EdGI
MdGI
MsGI
FvGI
AaGI
RcGI
RrGI
MiGI
JrGI
BpGI
MeGI
PtGIa
PtGIb
SvGI
GmGIc
GmGIa
GmGIb
MtGI
PsGI
AtGI
InGI
IbGI
SlGI
StGIa
StGIb
ZmGIb
ZmGIc
ZmGIa
OsGI
LpGI
BdGI
ScGI
TaGI
HvsGI
HvGI
MpGI
0.1

B

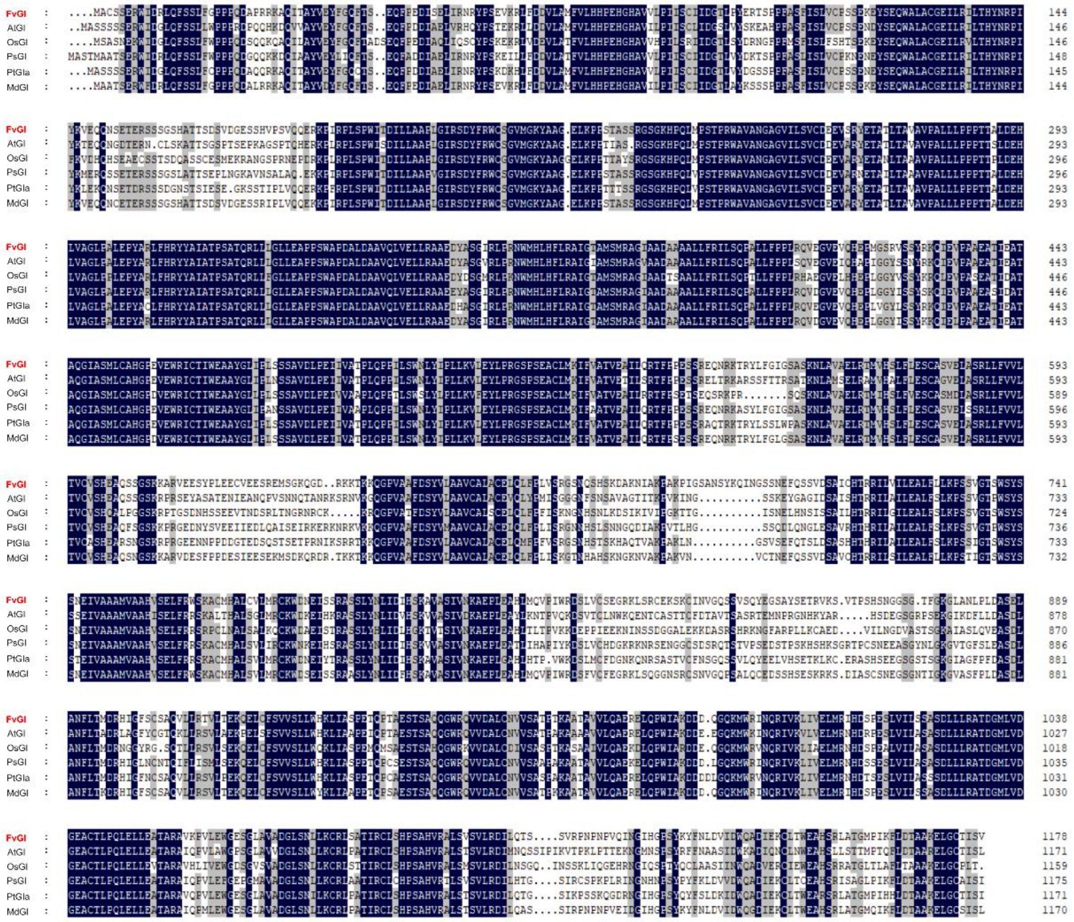
FvGI
AtGI
OsGI
PsGI
PtGIa
MdGI

**Fig. S1.** Phylogenetic analysis and multiple sequence alignment of GI homologs. (A) Phylogenetic analysis of GI proteins among different species. (B) Multiple sequence alignment of full-length GI protein sequences from species in which GI function has been characterized. Conserved amino acid residues are highlighted according to the degree of sequence conservation: black shading indicates positions with 100% conservation, grey represents ≥75% conservation, and white denotes ≤75% conservation. Species abbreviations: *Aa, Arabis alpina; Ag, Alnus glutinosa; At, Arabidopsis thaliana; Bd, Brachypodium distachyon; Bp, Betula platyphylla; Ed, Eriobotrya deflexa; Fv, Fragaria vesca; Gm, Glycine max; Hv, Hordeum vulgare; Ib, Ipomoea batatas; In, Ipomoea nil; Jr, Juglans regia; Lo, Lathyrus oleraceus; Lp, Lolium perenne; Md, Malus domestica; Me, Manihot esculenta; Mi, Mangifera indica; Mp, Marchantia polymorpha; Ms, Malus sylvestris; Mt, Medigago truncatula; Os, Oryza sativa; Pa, Prunus armeniaca; Pb, Pyrus x bretschneideri; Pd, Prunus dulcis; Pp, Prunus persica; Pt, Populus trichocarpa; Ra, Rubus arcutus; Rc, Rosa chinensis; Rr, Rosa rugosa; Sc, Secale cereale; Sl, Solanum lycopersicum; St, Solanum tuberosum; Sv, Salix viminalis; Zm, Ta, Triticum aestivum; Zea mays.* Details of protein sequences are found in Table S4.

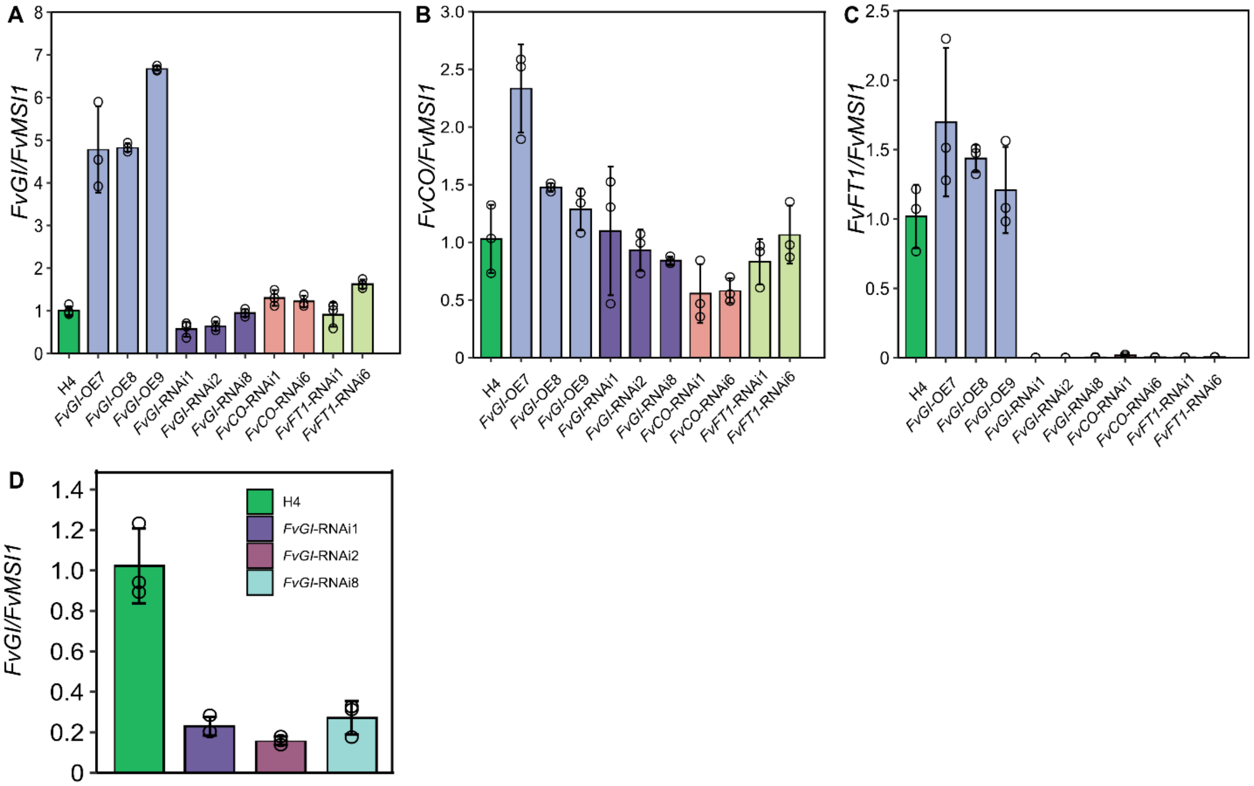

**Fig. S2**. Gene expression in H4 and transgenic lines. (A-C) RT-qPCR analysis *FvGI* (A), *FvCO* (B) and *FvFT1* (C) expression in leaf samples in H4, *FvGI*-OE lines and RNAi silencing lines for *FvGI*, *FvCO*, and *FvFT1*. Leaf samples including one leaflet per biological replicate (n = 3) were collected at ZT4 under $LD^{18h}$ conditions. Values are mean ± SD. (D) RT-qPCR analysis of *FvGI* in H4 and indicated RNAi lines. Leaf samples including one leaflet per biological replicate (n = 3) were collected at ZT8 under $LD^{18h}$ conditions. Values are mean ± SD.

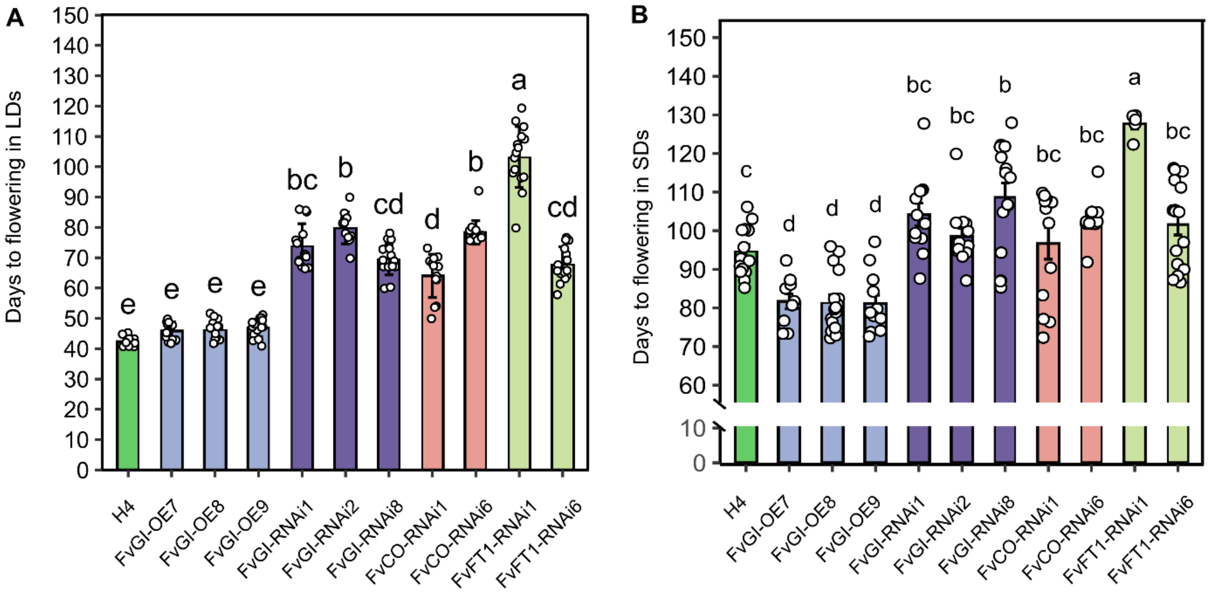


**Fig. S3**. Flowering time of H4 and transgenic lines. (A, B) Flowering time of indicated lines grown under $LD^{18h}$ (A) and $SD^{12h}$ (B) conditions. Different letters indicate statistically significant differences as determined by one-way ANOVA followed by Tukey's HSD multiple comparisons test ($P < 0.05$).

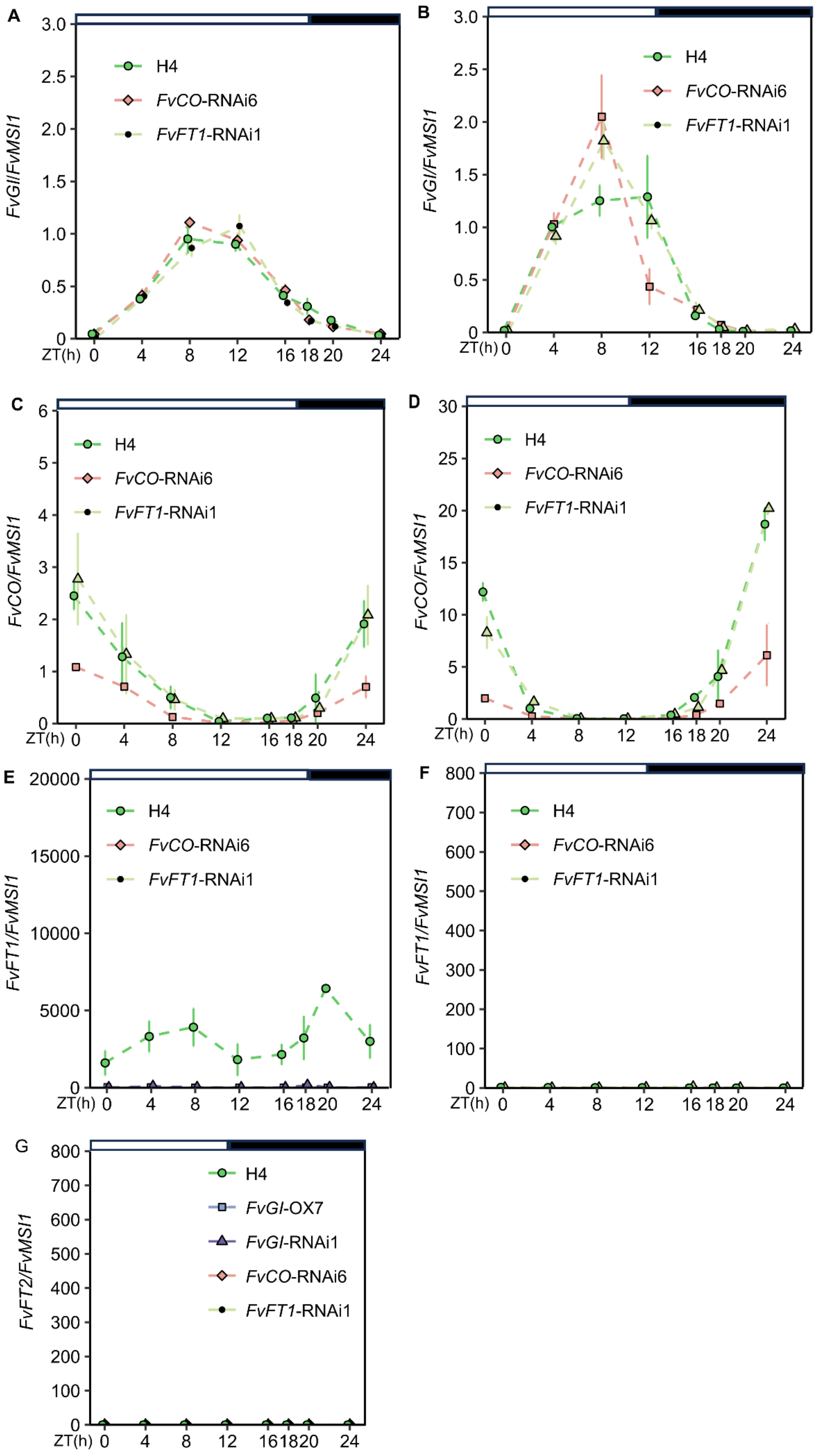
A
FvGI/FvMSI1
H4
FvCO-RNAi6
FvFT1-RNAi1
ZT(h)
B
FvGI/FvMSI1
C
FvCO/FvMSI1
D
FvCO/FvMSI1
E
FvFT1/FvMSI1
F
FvFT1/FvMSI1
G
FvFT2/FvMSI1
FvGI-OX7
FvGI-RNAi1

**Fig. S4**. Diurnal gene expression rhythms in H4 and transgenic lines. (A-F) Diurnal expression rhythms of *FvGI* (A, B), *FvCO* (C, D) and *FvFT1* (E, F) in leaves of H4, and *FvCO*- and *FvFT1*-RNAi lines grown under $LD^{18h}$ (left) and $SD^{12h}$ (right) conditions. (G) Diurnal gene expression analysis of *FvFT2* in leaf samples of H4 and its *FvGI*-OE and *FvGI*-, *FvCO*- and *FvFT1*- silencing lines grown under $SD^{12h}$ conditions. Values are mean ± SD (n = 3). White and black bars above the panels represent light and dark periods, respectively. ZT0 = lights on. Expression levels were measured in leaves after 3 weeks of light treatments using RT-qPCR.

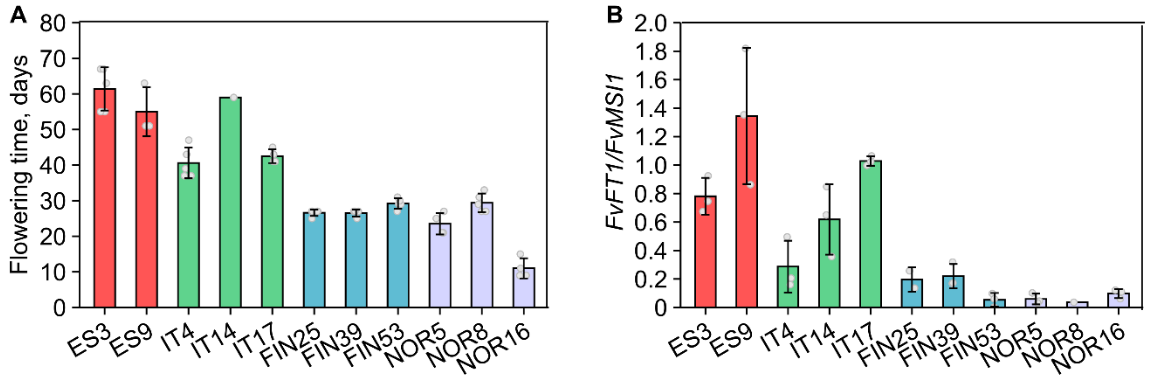


**Fig. S5**. Flowering time and *FvFT1* expression in natural accessions of *Fragaria vesca*. (A, B) Flowering time (A) and *FvFT1* expression levels (B) of indicated accessions. In A, plants (n = 5) were induced to flower under $SD^{12h}$ conditions (6 weeks), followed by flowering time observations under $LD^{18h}$. In B, leaf samples (n = 3) were collected at ZT4 under $LD^{18h}$ conditions. Values are mean ± SD.

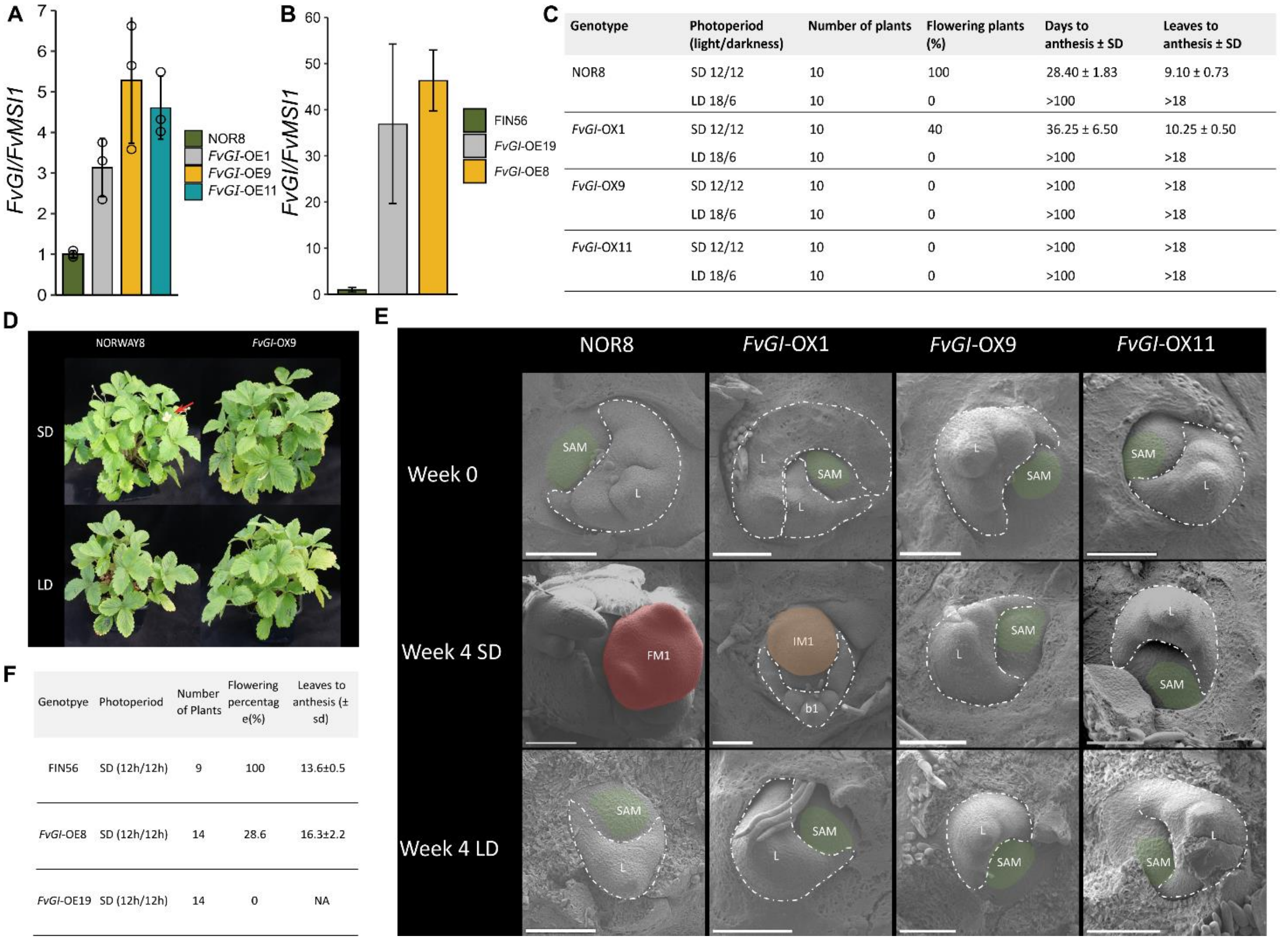


C

| Genotype | Photoperiod (light/darkness) | Number of plants | Flowering plants (%) | Days to anthesis ± SD | Leaves to anthesis ± SD |
|---|---|---|---|---|---|
| NOR8 | SD 12/12 | 10 | 100 | 28.40 ± 1.83 | 9.10 ± 0.73 |
| | LD 18/6 | 10 | 0 | >100 | >18 |
| *FvGI*-OX1 | SD 12/12 | 10 | 40 | 36.25 ± 6.50 | 10.25 ± 0.50 |
| | LD 18/6 | 10 | 0 | >100 | >18 |
| *FvGI*-OX9 | SD 12/12 | 10 | 0 | >100 | >18 |
| | LD 18/6 | 10 | 0 | >100 | >18 |
| *FvGI*-OX11 | SD 12/12 | 10 | 0 | >100 | >18 |
| | LD 18/6 | 10 | 0 | >100 | >18 |

F

| Genotpye | Photoperiod | Number of Plants | Flowering percentag e(%) | Leaves to anthesis (± sd) |
|---|---|---|---|---|
| FIN56 | SD (12h/12h) | 9 | 100 | 13.6±0.5 |
| *FvGI*-OE8 | SD (12h/12h) | 14 | 28.6 | 16.3±2.2 |
| *FvGI*-OE19 | SD (12h/12h) | 14 | 0 | NA |

**Fig. S6**. Gene expression and phenotypes of indicated *Fragaria vesca* genotypes. (A, B) RT-qPCR analysis *FvGI* in NOR8 and FIN56 accessions and indicated *FvGI* overexpression (OE) lines in shoot apex samples. For each biological replicate (n = 3), three shoot apices were harvested under $LD^{18h}$ conditions. Values are mean ± SD. (C) Flowering phenotypes of NOR8 and *FvGI*-OE lines. Flowering time is indicated as days to flowering and leaf numbers to flowering after indicated treatments of four weeks. After treatments, plants were transferred to $LD^{18h}$ for flowering time observations for up to 12 weeks. (D) Phenotypes of indicated treatments after photoperiodic treatments. Red arrow shows flower in NOR8 after $SD^{12h}$ conditions. (E) Scanning electron microscope images of meristems at week 0 and after 4 weeks under $LD^{18h}$ and $SD^{12h}$ treatments. All meristems remained vegetative under $LD^{18h}$ treatment. Under $SD^{12h}$ treatment, NOR8 displayed early stage of flower meristem (FM1), whereas a weaker *FvGI*-OE line had reached inflorescence meristem stage (IM1). Meristems of two strong *FvGI*-OE lines remained vegetative. White dashes: leaves. Green indicated area: vegetative shoot apical meristem. Orange indicated area: inflorescence meristem (IM1). Red area: flower meristem (FM1). Bars in all images indicate 100 µm. (F) Flowering time of FIN56 and its *FvGI*-OE lines. Plants were grown under SD$^{12h}$ conditions for five weeks, followed by flowering time observations under LD$^{18h}$.

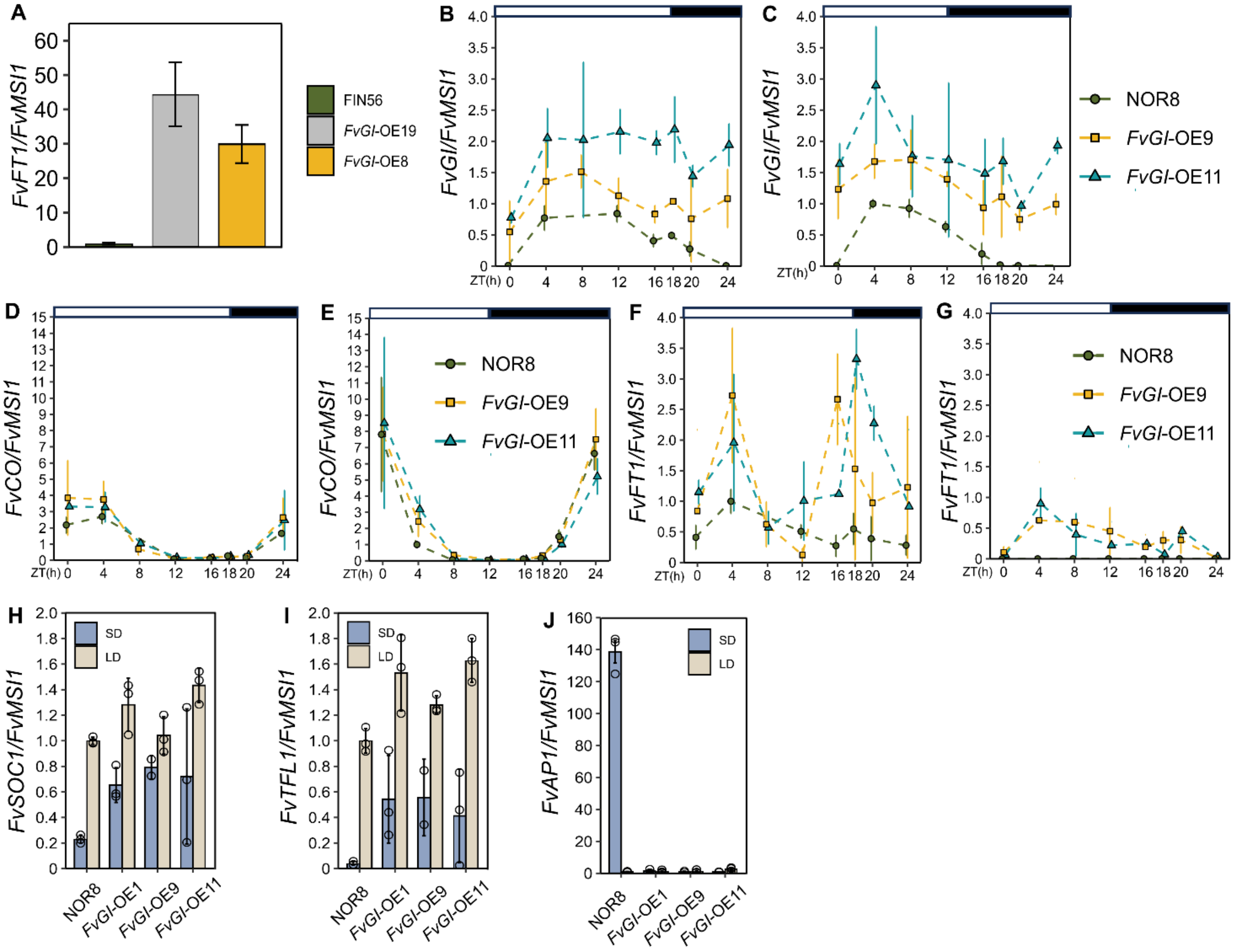


**Fig. S7**. Expression of flowering time genes in NOR8 and *FvGI* overexpression lines. (A) RT-qPCR analysis *FvFT1* in FIN56 and *FvGI*-OE lines in leaf samples collected at ZT4 under LD$^{18h}$ conditions. (B-G) The expression of *FvGI* (B, C), *FvCO* (D, E) and *FvFT1* (F, G) over 24 h diurnal cycle in NOR8 and *FvGI*-OE lines grown under LD$^{18h}$ (B, D, F) and SD$^{12h}$ (C, E, G) conditions for four weeks. Values are mean ± SD (n = 3). (H-J) RT-qPCR analysis *FvSOC1* (H), *FvTFL1* (I) and *FvAP1* (J) in shoot apex samples. Three shoot apices were harvested for each biological replicate after 4 weeks under indicated treatments. Values are mean ± SD (n = 3).

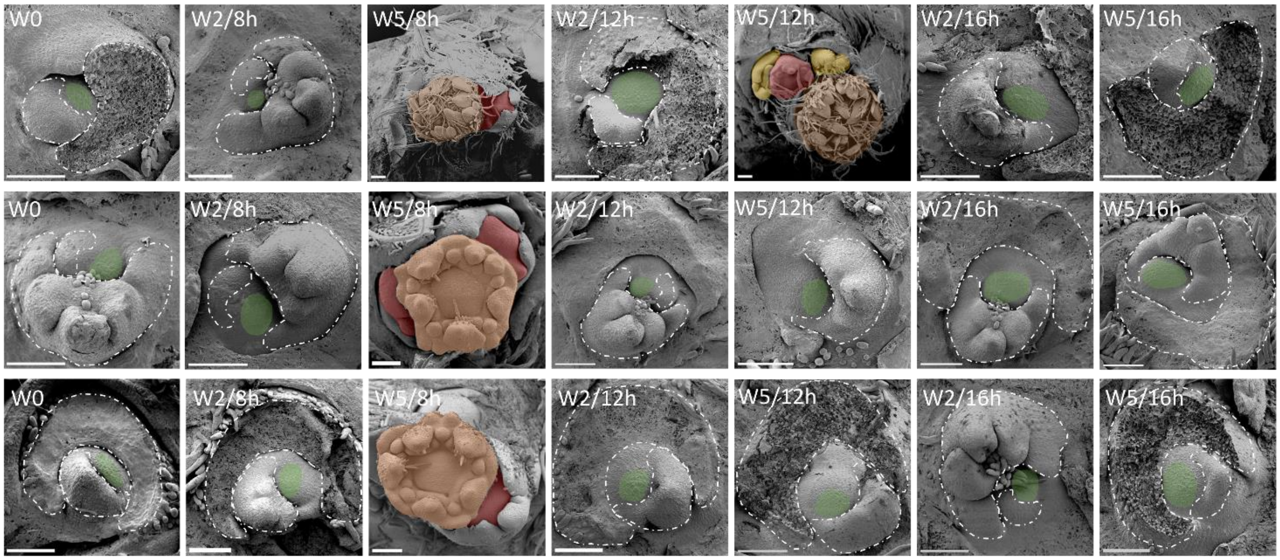


**Fig. S8.** Scanning electron microscope images of *Fragaria vesca* meristems. Shoot apical meristems (SAM) of NOR8 (row 1), and *FvGI* overexpression lines *FvGI*-OE9 (row 2) and *FvGI*-OE11 (row 3) after 0 (W0), 2 (W2) and 5 weeks (W5) under 8-h, 12-h, and 16-h photoperiods. After five weeks under 8-h photoperiod, NOR8 plants displayed late stage of flower meristem 1 (FM1, orange) and initiation of FM2 (red), whereas *FvGI*-OE plants exhibited an earlier FM1 (orange) stage with one or two visible later inflorescence meristems (IM2, red). After five weeks under 12-h photoperiod, more advanced developmental stage of NOR8 was found compared with NOR8 under 8-h photoperiod, with visible FM1 (orange), FM2 (red) and two lateral IM3 (yellow) in the shoot apex. Vegetative SAMs (green) were found in *FvGI*-OE plants under 12-h photoperiod and in all plants under 16-h photoperiod. White dashes area indicates leaves. Scale bars in all images are equal to 100 $\mu$m.

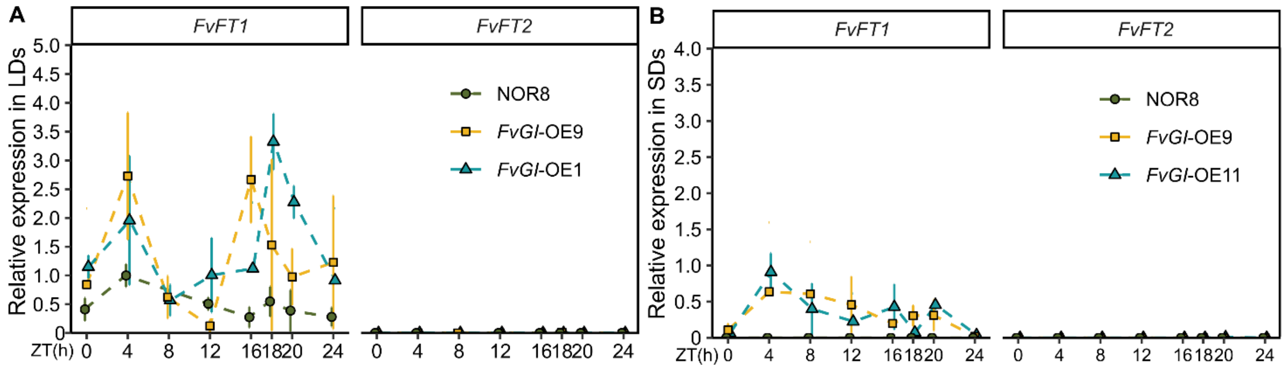


**Fig. S9**. Expression of *FvFT1* and *FvFT2* in NOR8 and *FvGI* overexpression lines. (A, B) Gene expression over 24-h cycle was analyzed in leaves of NOR8 and its *FvGI*-OE lines harvested under $LD^{18h}$ (A) and $SD^{12h}$ (B) conditions. Values are mean ± SD (n = 3).

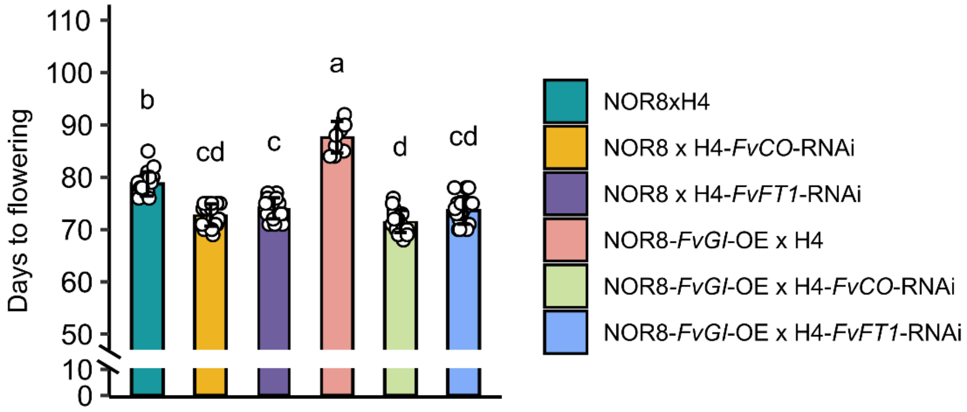


**Fig. S10.** Flowering time of indicated hybrids. Four-week-old plants, clonally propagated from hybrid seedlings were subjected to $SD^{12h}$ at 17°C for eight weeks followed by phenotyping under $LD^{18h}$. Values are mean ± SD (n = 18). Different letters indicate statistically significant differences as determined by one-way ANOVA followed by Tukey's HSD multiple comparisons test ($P < 0.05$).

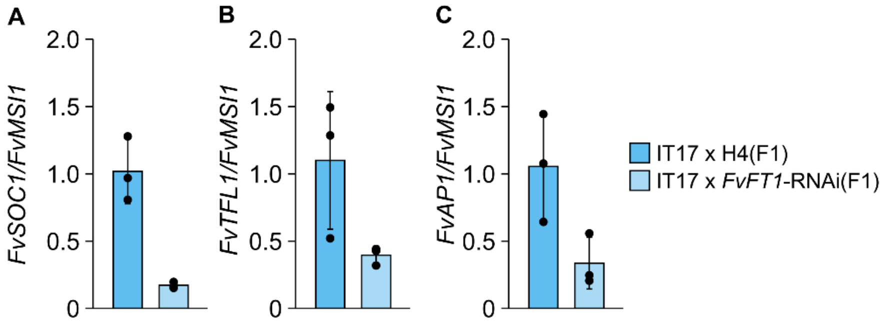


**Fig. S11.** Expression of flowering genes in indicated hybrids. (A-C) The expression of *FvSOC1* (A), *FvTFL1* (B) and *FvAP1* (C) in shoot apices harvested under $LD^{18h}$ conditions in IT17 x H4 and IT17 x H4 *FvFT1*-RNAi F1 plants. Three shoot apices were harvested for each biological replicate. Values are mean ± SD (n = 3).

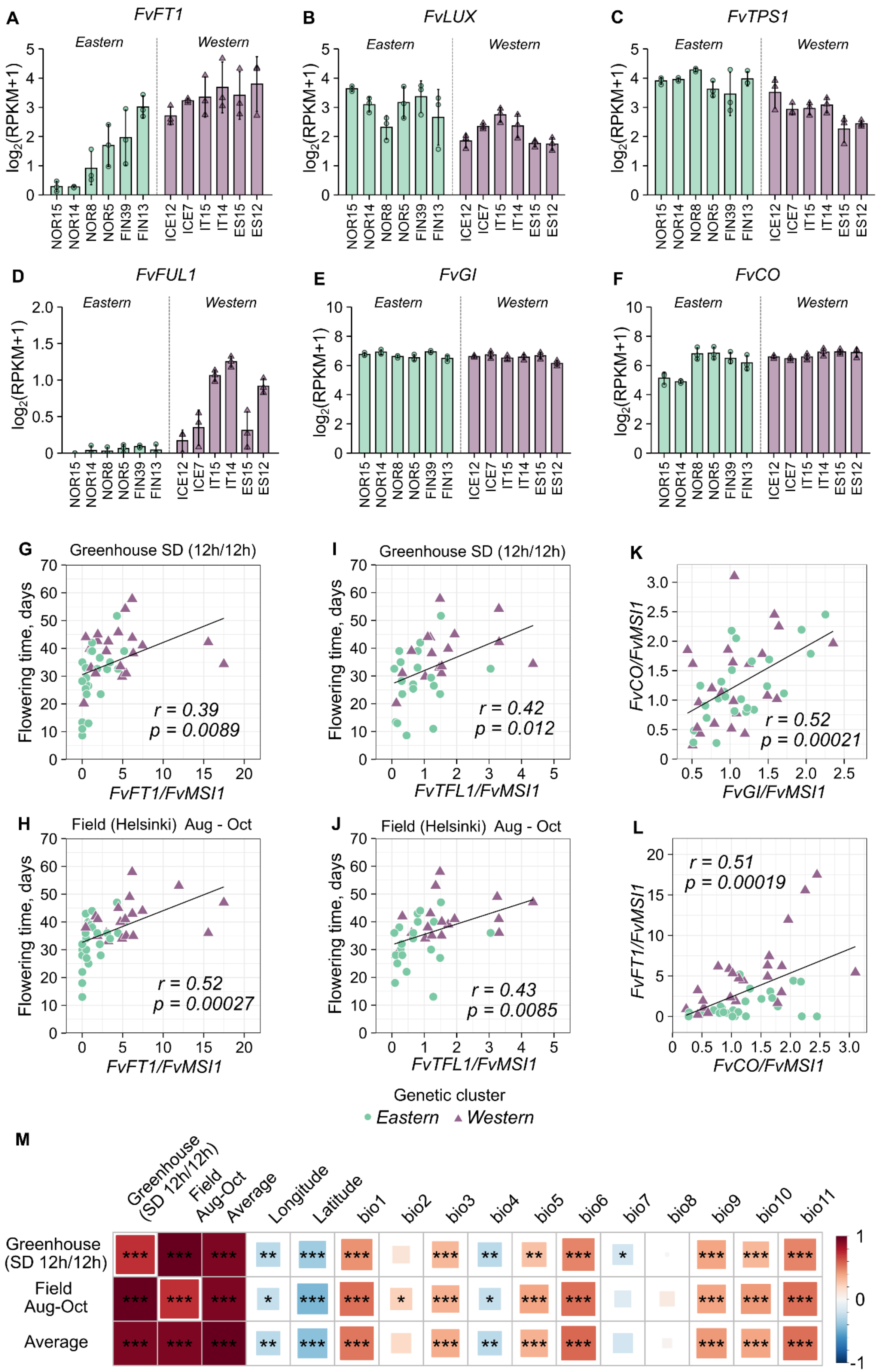
A
FvFT1
B
FvLUX
C
FvTPS1
D
FvFUL1
E
FvGI
F
FvCO
Eastern
Western
log2(RPKM+1)
NOR15
NOR14
NOR8
NOR5
FIN39
FIN13
ICE12
ICE7
IT15
IT14
ES15
ES12
G
Greenhouse SD (12h/12h)
Flowering time, days
r = 0.39
p = 0.0089
FvFT1/FvMSI1
I
Greenhouse SD (12h/12h)
r = 0.42
p = 0.012
FvTFL1/FvMSI1
K
FvCO/FvMSI1
r = 0.52
p = 0.00021
FvGI/FvMSI1
H
Field (Helsinki) Aug - Oct
r = 0.52
p = 0.00027
FvFT1/FvMSI1
J
Field (Helsinki) Aug - Oct
r = 0.43
p = 0.0085
FvTFL1/FvMSI1
L
r = 0.51
p = 0.00019
FvFT1/FvMSI1
FvCO/FvMSI1
Genetic cluster
Eastern
Western
M
Greenhouse (SD 12h/12h)
Field Aug-Oct
Average
Longitude
Latitude
bio1
bio2
bio3
bio4
bio5
bio6
bio7
bio8
bio9
bio10
bio11
Greenhouse (SD 12h/12h)
Field Aug-Oct
Average
1
0
-1

**Fig. S12.** Comparison of gene expression levels in *Fragaria vesca* accessions and correlations between gene expression, flowering time, and bioclimatic and geographical variables. (A-F) Expression levels of *FvFT1* (A), *FvLUX* (B), *FvTPS1* (C), *FvFUL1* (D), *FvGI* (E) and *FvCO* (F) in six eastern and six western accessions. Leaf samples were collected at ZT4 under $LD^{18h}$ conditions and analysed using RNA sequencing. (G-J) Pearson's correlations between relative expression of *FvFT1* (G and H) in leaves ($LD^{18h}$, ZT16) and *FvTFL1* (I and J) in shoot apices ($LD^{18h}$) and flowering time observations in a greenhouse after $SD^{12h}$ treatment of six weeks (G and I) or in plants induced to flower in the field in Helsinki in August-October (H and J). After treatments, flowering time observations were carried out in the greenhouse under $LD^{18h}$. The gene expression data is from Fig. 5h. (K) Pearson's correlation between relative expression of *FvGI* and *FvCO*. (L) Pearson's correlation between relative expression of *FvCO* and *FvFT1*. In K and L, leaf samples used in Fig. 5e were analyzed. (M) Heatmap showing Pearson's correlations between flowering time and temperature related bioclimatic variables, longitude and latitude. Flowering time data from $SD^{12h}$ treatment (179 accessions), field experiment (181 accessions), and their average (179 accessions) were used. Asterisks mark the significance levels (***, $p < 0.001$; **, $p < 0.01$; *, $p < 0.05$, Bonferroni adjusted).
